# PharmOmics: A Species- and Tissue-specific Drug Signature Database and Online Tool for Drug Repurposing

**DOI:** 10.1101/837773

**Authors:** Yen-Wei Chen, Graciel Diamante, Jessica Ding, Thien Xuan Nghiem, Jessica Yang, Sung-min Ha, Peter Cohn, Douglas Arneson, Montgomery Blencowe, Jennifer Garcia, Nima Zaghari, Paul Patel, Xia Yang

## Abstract

Drug development has been hampered by a high failure rate in clinical trials due to efficacy or safety issues not predicted by preclinical studies in model systems. A key contributor is our incomplete understanding of drug functions across organ systems and species. Therefore, elucidating species- and tissue-specific actions of drugs can provide systems level insights into therapeutic efficacy, potential adverse effects, and interspecies differences that are necessary for more effective translational medicine. Here, we present a comprehensive drug knowledgebase and analytical tool, PharmOmics, comprised of genomic footprints of drugs in individual tissues from human, mouse, and rat transcriptome data from GEO, ArrayExpress, TG-GATEs, and DrugMatrix. Using multi-species and multi-tissue gene expression signatures as molecular indicators of drug functions, we developed gene network-based approaches for drug repositioning. We demonstrate the potential of PharmOmics to predict drugs for new disease indications and validated two predicted drugs for non-alcoholic fatty liver disease in mice. We also examined the potential of PharmOmics to identify drugs related to hepatoxicity and nephrotoxicity. By combining tissue- and species-specific *in vivo* drug signatures with biological networks, PharmOmics serves as a complementary tool to support drug characterization.

## Background

Drug development has been challenging and costly over the past decades due to the high failure rate in clinical trials (1). Most drugs with excellent efficacy and safety profiles in preclinical studies often encounter suboptimal efficacy or safety concerns in humans (2). This translational gap is likely attributable to our incomplete understanding of the systems level activities of drugs in individual tissues and organ systems (3) as well as the differences between humans and model systems (4).

Drug activities can be captured by gene expression patterns, commonly referred to as gene signatures. By measuring how a pharmacological agent affects the gene signature of a cell or tissue type in a particular species, we can infer the cell- or tissue-specific biological pathways involved in therapeutic processes or toxicological responses. This concept has prompted drug repositioning studies and provided important predictions for repurposing approved drugs for new disease indications (5–10). Similarly, gene signatures can reveal mechanisms underlying adverse drug reactions (ADRs) and be leveraged to predict ADRs as previously shown for liver and kidney toxicity (11–13).

A drug may affect different molecular processes between tissues, providing treatment effects in the desired target tissue(s) but causing toxicity or ADRs in other tissues. Therefore, tissue-specific drug signatures will offer a more systematic understanding of drug actions *in vivo*. In addition, rodent models have been commonly used in toxicology and preclinical studies, yet species-specific effects of drugs have been observed (14) and underlie the lack of efficacy or unexpected ADRs of certain drugs when used in humans (15). Therefore, understanding the species-specific molecular effects of drugs is of high biological importance. A detailed species- and tissue-specific drug genomic signature database will significantly improve our understanding of the molecular networks affected by drugs and facilitate network-based drug discovery and ADR prediction for translational medicine.

The potential of using gene signatures to facilitate target and toxicity identification has led to several major efforts in characterizing genomic signatures related to drug treatment (8,16–18). However, none of the existing platforms offer comprehensive cross-tissue and cross-species *in vivo* assessments of drug activities to allow predictions of drug effects on individual tissues and to help assess the translational potential of a drug based on consistencies or discrepancies between species. For instance, the comparative toxicogenomics database (CTD), a literature-based resource curating chemical-to-gene/protein associations as well as chemical-to-disease and gene/protein-to-disease connections (16), lacks the cellular and tissue context of the curated interactions. More systematic, data-driven databases like CMAP (8) and LINC1000 (17) focus on characterizing and cataloging the genomic footprints of more than ten thousand chemicals using *in vitro* cell lines (primarily cancer cell lines) to offer global views of molecular responses to drugs in individual cellular systems. However, these *in vitro* cell-line based gene signatures may not always capture *in vivo* tissue-specificity of drug activities. To move into *in vivo* systems, large toxicogenomics databases like TG-GATEs (19) and DrugMatrix from the National Toxicology Programs of the National Institute of Environmental Health Sciences (https://ntp.niehs.nih.gov/drugmatrix/index.html) have become available to provide unbiased transcriptome assessment for heart, muscle, liver, and kidney systems. However, information about other organ systems is limited. Efforts to analyze publicly deposited transcriptomic datasets in GEO (20) and ArrayExpress (21), which have broader tissue coverage, for individual drugs have been described (18), but systematic annotation and integration of species- and tissue-specific effects of drugs have not been achieved.

Here, we present a database that contains 13,382 rat, human, and mouse transcriptomic datasets across >20 tissues covering 941 drugs. We evaluated the tissue- and species-specificity of drug signatures as well as the performance of these signatures in gene network-based drug repositioning, toxicity prediction, target identification, and comparisons of molecular activities between tissues and species. The drug signatures are hosted on an interactive web server, PharmOmics, to enable public access to drug signatures and integrative analyses for drug repositioning (http://mergeomics.research.idre.ucla.edu/runpharmomics.php).

## Methods

### Curation of tissue- and species-specific drug transcriptomic datasets

As illustrated in **Figure 1**, we compiled a list of clinically relevant drugs, including 766 FDA approved drugs from Kyoto Encyclopedia of Genes and Genomes (KEGG) (16), which overlapped with drugs from the US Food and Drug Administration (FDA), European Medical Agency, and Japanese Pharmaceuticals and Medical Devices Agency, with an additional 175 chemicals from TG-GATEs (19) and DrugMatrix (https://ntp.niehs.nih.gov/drugmatrix/index.html). The compiled drug list was queried against GEO, ArrayExpress, TG-GATEs, and DrugMatrix to identify datasets as of January 2018. Duplicated datasets between data repositories were removed. We developed a semi-automated pipeline combining automated search with manual checking to identify relevant datasets for drug treatment. The automated process first extracts datasets containing drug generic names or abbreviations and then inspects the potential datasets for availability of both drug treatment and control labels in the constituent samples. We also manually checked the recorded labels identified by the automated process to validate the labels and remove potential false detections. Only datasets with n>=3/group in both drug treatment and control groups were included in our downstream analyses. Although a larger sample size is desired, the majority (78.7%) of drug transcriptome datasets have n=3/group, 20.9% datasets have n=2/group, and <1% datasets have n>3/group (**Supplementary Table 1**).

**Figure 1.**
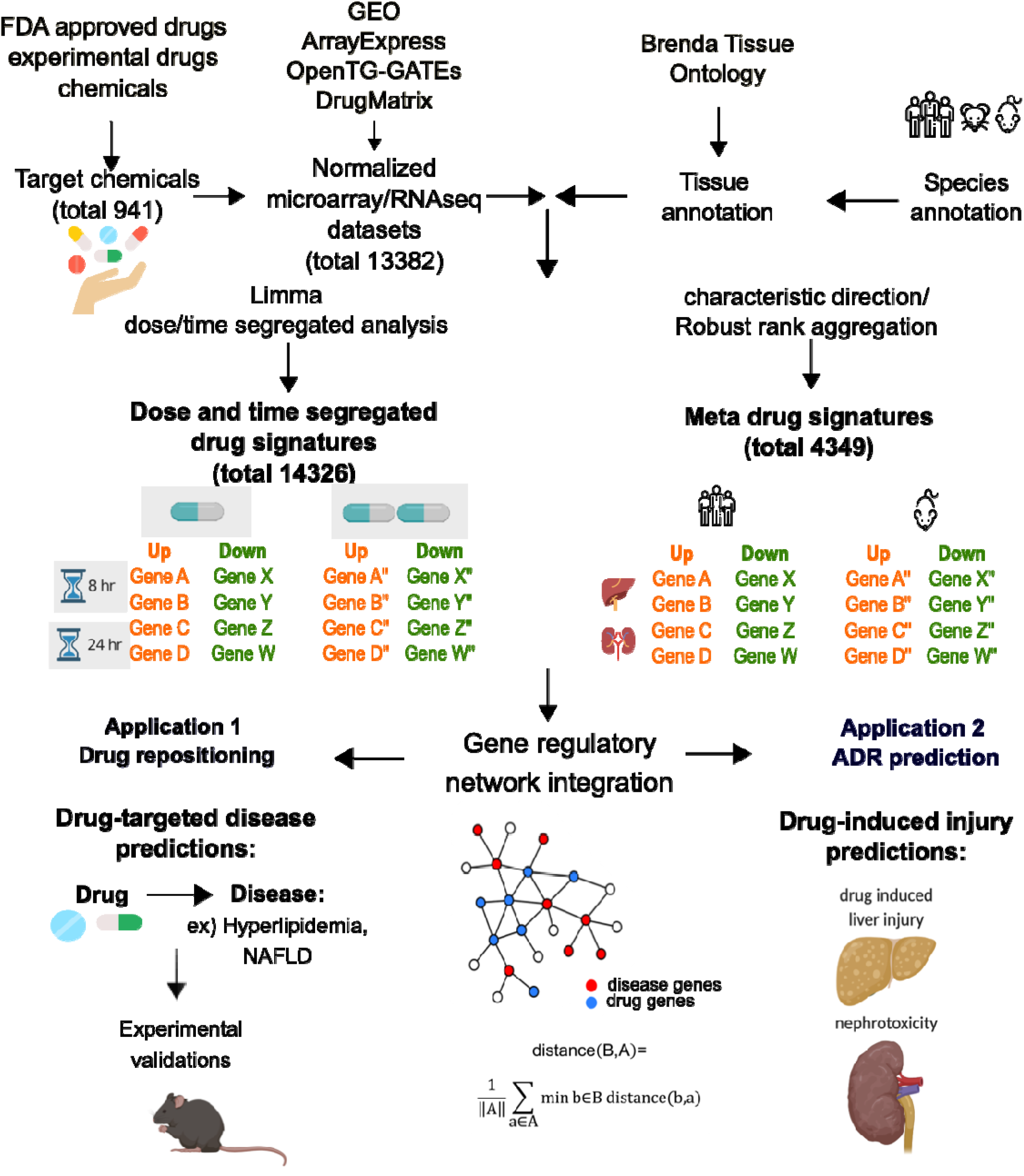
PharmOmics data processing pipeline. FDA approved drugs based on KEGG database were searched against GEO, ArrayExpress, TG-GATEs, and DrugMatrix data repositories. Additional experimental drugs and chemicals from TG-GATEs and DrugMatrix were also included. Only datasets with drug treatment and control samples were retrieved. Datasets were first annotated with tissue and species information, followed by retrieval of dose/time-segregated or meta-analysis drug signatures using two different methods. Dose/time-segregated signatures were retrieved from individual datasets using LIMMA. Meta signatures across datasets of the same drugs were obtained by first applying GeoDE to obtain a ranked gene list for each treatment experiment, followed by meta-analysis using the Robust Rank Aggregation method. These signatures were used to conduct drug repositioning analysis and hepatotoxicity/nephrotoxicity prediction based on direct gene overlaps or a gene network-based approach.

### Obtaining drug treatment signatures stratified by species and tissues

Species and tissue labels were retrieved based on the metadata of each dataset. Tissue names were standardized based on the Brenda Tissue Ontology (22). We implemented a search procedure to climb the ontology tree structure to determine the suitable tissue annotations. This was done by first building a priority list of widely used tissues/organs in toxicological research using the Brenda Tissue Ontology tree system. Tissue/organ priority order was set to “kidney”, “liver”, “pancreas”, “breast”, “ovary”, “adipose tissue”, “cardiovascular system”, “nervous system”, “respiratory system”, “urogenital system”, “immune system”, “hematopoietic system”, “skeletal system”, “integument" (endothelial and skin tissue), “connective tissue”, “muscular system”, “gland”, “gastrointestinal system”, and “viscus” (other non-classified tissue). Tissue terms relevant to each of these tissues or organs were curated from the ontology tree into a tissue/organ ontology table. Next, we looked up terms from our tissue/organ ontology table in the Cell/Organ/Tissue column of the metadata in each transcriptomic dataset. If a tissue/organ term was not found, we searched the title and summary columns of the metadata as well to retrieve additional information. When the search returned multiple tissue terms (for example, breast cancer cell line may be categorized as both epithelial and breast organ), we used the term with the highest priority as described above. We prioritized the tissue terms based on the relevance to toxicology to make tissue assignments unique for each dataset to reduce ambiguity. Manual checking was conducted to confirm the tissue annotation for each dataset.

For each gene expression dataset, normalized data were retrieved, and quantile distribution of the values were assessed. When a dataset was not normally distributed, log2-transformation using GEO2R (20) was applied. To identify differentially expressed genes (DEGs) representing drug signatures, two different strategies were used. First, the widely used DEG analysis method LIMMA (23) was applied to obtain dose and time segregated signatures under FDR < 0.05. To overcome the low sample size issue and obtain “consensus” drug signatures for a drug/chemical, LIMMA was also applied to datasets where multiple doses and treatment durations were tested, and treatment effect were derived by combining dose/time experiments for the same drug/chemical in each study. Second, we leveraged different studies for the same drugs or chemicals in the same tissue and species to derive meta-analysis signatures. To address heterogeneity in study design, platforms, sample size, and normalization methods across different data repositories, we applied the characteristic direction method from the GeoDE package (24) to derive consistent DEGs for each drug across different data sources. GeoDE was designed to accommodate heterogenous datasets that have differing parameters and outputs between treatment and control groups. It uses a “characteristic direction” measure to identify biologically relevant genes and pathways. The normalized characteristic directions for all genes were then transformed into a non-parametric rank representation. Subsequently, gene ranks of a particular drug from the same tissue/organ system and the same organism were aggregated across datasets using the Robust Rank Aggregation method (25), a statistically controlled process to identify drug DEGs within each tissue for each species. Robust Rank Aggregation provides a non-parametric meta-analysis across different ranked lists to obtain commonly shared genes across datasets, which avoids statistical issues associated with heterogeneous datasets. It computes a null distribution based on randomized gene ranks and then compares the null distribution with the empirical gene ranks to obtain a p-value for each gene. The robust rank aggregation process was done for the upregulated and downregulated genes separately to obtain DEGs for both directions under Bonferroni-adjusted p-value < 0.01, a cutoff implemented in the Robust Rank Aggregation algorithm. To obtain species-level signatures for each drug, we further aggregated DEGs across different organs tested for each drug within each species.

Pathway analysis of individual drug signatures was conducted using Enrichr (26) by intersecting each signature with pathways or gene sets from KEGG (27) and gene ontology biological process (GOBP) terms (28). In addition, pathway enrichment analysis based on network topology analysis (29) was conducted using ROntoTools (30). Pathways at false discovery rate (FDR) < 0.05 were considered significant in both methods.

We curated 14,366 drug signatures segregated by treatment dosage and duration, tissue, and species, covering 719 drugs and chemicals, among which 544 are FDA approved. In addition, our meta signatures is a consensus of 4,349 signatures covering 551 drugs across treatment regimens. In total, the entire database is based on 13,382 rat, human, and mouse transcriptomic datasets across >20 tissue or organ systems across 941 drugs and chemicals from GEO, ArrayExpress, DrugMatrix, and TG-GATEs to derive drug signatures. These rat, human, and mouse datasets cover >20 tissue or organ systems. The toxicogenomics databases TG-GATEs and DrugMatrix mainly contain liver and kidney datasets from rats, while public data repositories GEO and ArrayExpress contain datasets with broader tissue and species coverage (**Figure 2A**). Overall, the rat datasets are mainly from liver and kidney whereas human and mouse datasets also contained signatures from other tissues and organs such as breast and the nervous system (**Figure 2B**). There is also a species bias between the data repositories; GEO covered more mouse and human datasets, TG-GATEs mainly has human and rat datasets, and DrugMatrix curated more rat datasets (**Figure 2C)**.

**Figure 2.**
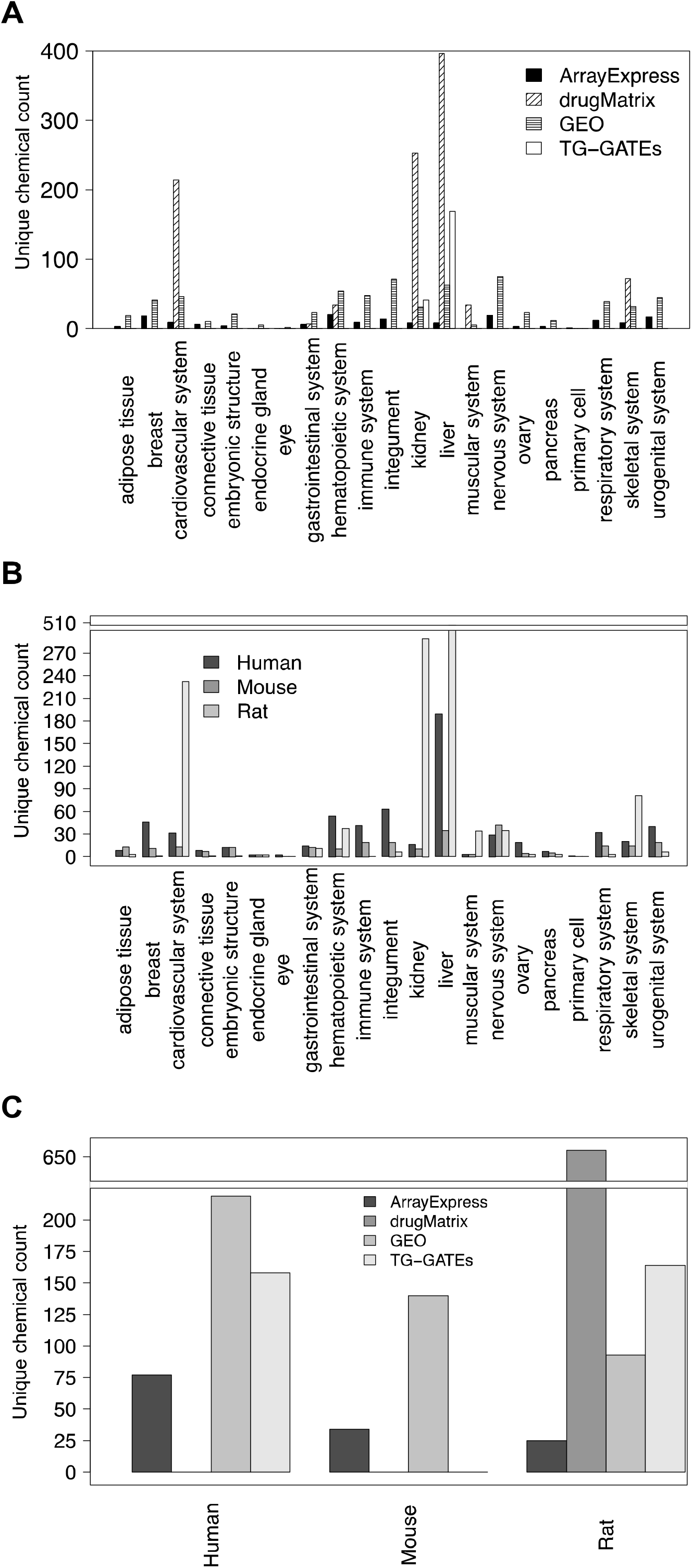
Summary of available datasets based on data sources, tissues, and species. Y-axis indicates unique dataset counts, and X-axis indicates (A) tissue and data resources, (B) tissue and species, and (C) data resources and species.

### Comparison of PharmOmics with existing drug signature platforms

To assess the degree of agreement in drug signatures between the PharmOmics database and existing platforms, we compared PharmOmics with the CREEDS (18) and L1000FWD (31) databases, for which drug signatures are accessible **(Supplementary methods)**. As shown in **Supplementary Figure 1**, both the PharmOmics dose/time-segregated signatures and the meta signatures showed better concordance with the two existing platforms than the agreement between CREEDS and L1000FWD, as reflected by higher overlap fold enrichment score and lower statistical p values. The three platforms have differences in the datasets and analytical strategies and therefore are complementary. Due to the lack of full access to CMAP signatures, we were not able to systematically compare PharmOmics against CMAP.

### Web server implementation of PharmOmics

To allow easy data access and use of PharmOmics, we have created a freely accessible web tool deployed on the same Apache server used to host Mergeomics (32), a computational pipeline for integrative analysis of multi-omics datasets to derive disease-associated pathways, networks, and network regulators (http://mergeomics.research.idre.ucla.edu).

The PharmOmics web server features three functions (**Figure 3**). First, it allows queries for species- and tissue-stratified drug signatures and pathways for both the dose/time-segregated and meta signatures. Details of statistical methods (e.g, LIMMA vs characteristic direction), signature type (dose/time-segregated vs meta), and datasets used are annotated. The drug query also includes a function for DEG and pathway signature comparisons between user-selected species and tissues which can be visualized and downloaded. Second, it features a network drug repositioning tool that is based on the connectivity of drug signatures in PharmOmics to user input genes such as a disease signature. This tool requires a list of genes and a gene network that can be chosen from our preloaded gene regulatory networks if relevant or a custom upload (see Applications below for details in implementation). In the output, Z-score and p-value results of network repositioning are displayed and available for download. In addition, we list the overlapping genes between drug signatures in the given network and the input genes, the drug genes with direct connections to input genes through one-edge extension, and input genes with one-edge connections to drug genes in the downloadable results file. The output page also provides network visualization which details the genes affected by a drug and their overlap with and direct connections to user input genes using Cytoscape.js. The network nodes and edges files are also available for download and can be used on Cytoscape Desktop. **Figure 4** shows the web interface of the input submission form (**Figure 4A**) and results display of the network repositioning tool using a sample liver network and a sample hyperlipidemia gene set (**Figure 4B**). Lastly, the web server offers a gene overlap-based drug repositioning tool that assesses direct overlap between drug gene signatures and user input genes. Gene overlap-based drug repositioning requires a single list of genes or separate lists of upregulated and downregulated genes and outputs the Jaccard score, odds ratio, Fisher’s exact test p-value, within-species rank, and gene overlaps for drugs showing matching genes with the input genes. This gene overlap-based approach is similar to what was implemented in other drug repositioning tools, but the network-based repositioning approach is unique to PharmOmics.

**Figure 3.**
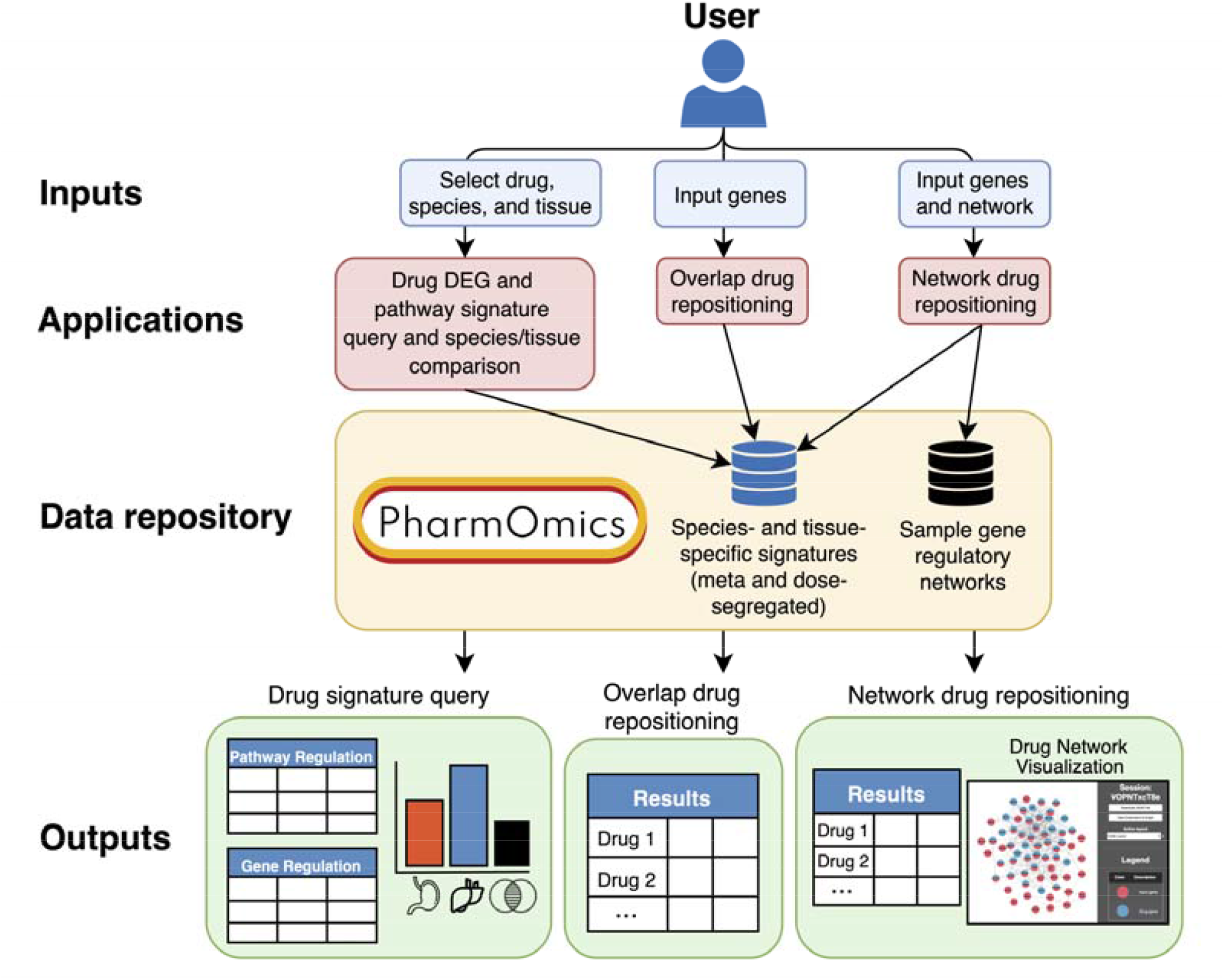
PharmOmics web server. The web server hosts drug signature and pathway queries, between-tissue and between-species drug signature comparisons, and network-based and gene overlap-based drug repositioning. Users are able to query, download, and perform drug repositioning using all species- and tissue-specific meta and dose/time-segregated signatures. Interactive results tables and network visualizations are displayed on the website and available for download.

**Figure 4.**
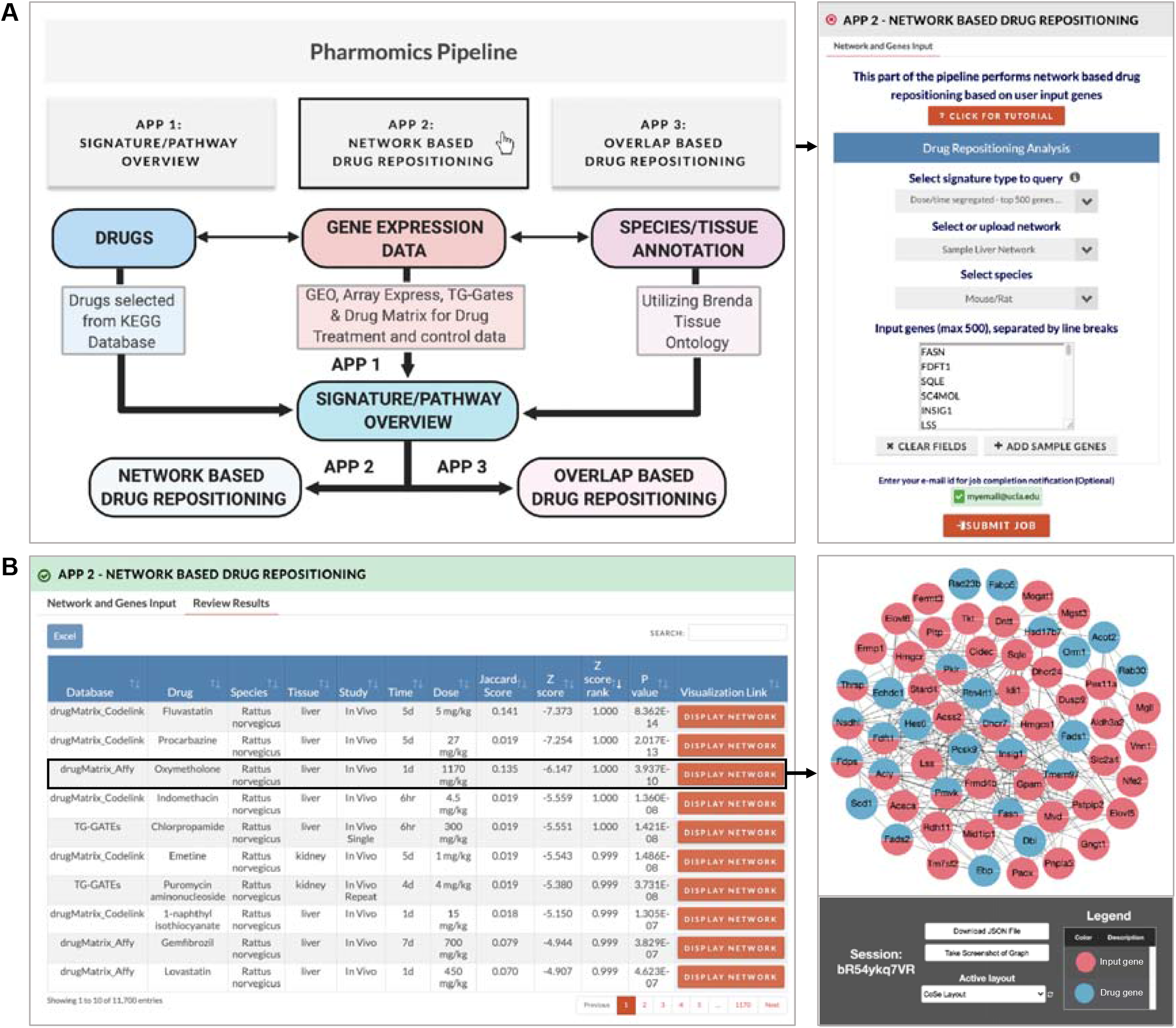
User interface of network drug repositioning web tool using sample hyperlipidemia gene set and sample mouse Bayesian gene regulatory network. (A) Inputs to network drug repositioning includes i) signature type to query (meta-analyzed, dose/time-segregated with top 500 genes per signature, or dose/time-segregated with all genes), ii) network (custom upload or select a sample network), iii) species (relating to the species of the network being used), and iv) genes. In this case we choose dose/time-segregated signatures using top 500 genes, a sample liver network, mouse/rat species, and the sample hyperlipidemia gene set (loaded from ‘Add sample genes’). If human gene symbols are provided with the ‘Mouse/Rat’ species selection, the genes will be converted to mouse/rat symbols. (B) After the job is complete, the results file is displayed on the website and available for download. A subset of the drug network containing the drug genes that are first neighbors to input genes and all input genes can be visualized using the “Display Network” button which will load an interactive display of the subnetwork topology. The oxymetholone drug signature in rat liver is a top hit, and the drug network is shown on the right. Additional data in the downloadable results file include the genes that are both a drug gene and an input gene in the network, drug genes that are directly connected (first neighbor) to input genes, and input genes directly connected to drug genes.

### Experimental methods for NAFLD drug validation

Seven-week old C57BL/6 male mice were purchased from the Jackson Laboratory (Bar Harbor, ME). After acclimation the animals were randomly assigned to four experimental groups (n=7-9/group) on different diets/treatments: regular chow diet (Control) (Lab Rodent Diet 5053, St. Louis, MO), high fat high sucrose (HFHS) diet (Research Diets-D12266B, New Brunswick, NJ) to induce hepatic steatosis, a key NAFLD phenotype, HFHS diet with fluvastatin treatment (NAFLD + Flu), and HFHS diet with aspirin treatment (NAFLD + Asp). The target intake concentrations of fluvastatin and aspirin were 15mg/kg and 80 mg/kg, respectively, which were chosen based on doses used in previous studies that did not show toxicity (33,34). These experimental diets were then administered for 10 weeks. The average fluvastatin intake was 14.98 mg/kg/day, and the average aspirin intake was 79.67 mg/kg/day.

During drug treatment, metabolic phenotypes such as body weight, body fat and lean mass composition were monitored weekly. Fat and lean mass were measured with Nuclear Magnetic Resonance (NMR) Bruker minispec series mq10 machine (Bruker BioSpin, Freemont, CA). For metabolic phenotypes measured at multiple time points (body weight gain and adiposity), differences between groups were analyzed using a 2-way ANOVA followed by Sidak’s multiple comparisons test. At the end of treatment, livers from all groups were weighed, flash frozen, and stored at −80◻°C until lipid analysis. Hepatic lipids were extracted using the Folch method as previously described (35). The lipid extracts were analyzed by the UCLA GTM Mouse Transfer Core for triglyceride (TG), total cholesterol (TC), unesterified cholesterol (UC), and phospholipids (PL) levels by colorimetric assay from Sigma (St. Louis, MO) according to the manufacturer’s instructions. All animal experiments were approved by the UCLA Animal Research Committee.

## Results

### Evaluating the ability of PharmOmics to extract drug targets and target pathways

It remains unclear whether drug DEGs reflect drug targets. To evaluate this possibility, we retrieved known targets for the drugs included in PharmOmics from the DrugBank database (36) and used three different methods to evaluate the potential of DEGs for drug target identification. The first method assessed direct gene overlaps between known drug targets and DEG signatures. The second assessed overlaps between known drug target pathways and drug DEG pathways from pathway enrichment analysis. The last method was based on whether known drug targets were within the close neighborhood of drug DEGs in molecular networks, including the STRING network (37) and tissue-specific Bayesian networks (BNs) **(Supplementary methods)**. For drugs with multiple dose and time regimens, only the signature with the best performance was used in these analyses.

The drug target recovery rates using PharmOmics drug DEGs for gene overlap, pathway overlap, STRING network overlap, and liver BN overlap with liver DEGs were 22%, 59.1%, 41.7%, and 60.2%, respectively, and were significantly higher than the rates using random genes (**Supplementary Table 2)**. Although these rates are low, gene overlap drug target recovery rate using PharmOmics signatures was higher than using CMAP (14%) and L1000 (17%) signatures, and drug target recovery was improved by pathway and network approaches. Notably, matching the tissue between DEGs and network improved the target detection rate. However, we note that while the pathway- and network-based approaches increased the detection rate for true drug targets, the number of false positives was also increased. Overall, our results show that although PharmOmics has certain value in drug target and pathway retrieval as shown by better performance than random genes and other platforms, the retrieval rate is low. These results suggest that DEGs do not recover direct drug targets well but more likely reflect target-related pathways, and caution should be taken when using DEGs for target identification.

### Utility of PharmOmics drug signatures in retrieving known therapeutic drugs for various diseases

We next evaluate the ability of PharmOmics drug signatures to identify drugs for diseases based on overlaps or network connectivity in gene signatures matched by tissue. We hypothesized that if a drug is useful for treating a disease, the drug signatures and disease signatures likely target similar pathways and therefore have direct gene overlaps or connect extensively in gene networks. For gene overlap-based drug repositioning, we calculate the Jaccard score, gene overlap fold enrichment, and Fisher’s exact test p values as the overlap measurements. For network-based drug repositioning, we used a network proximity measurement between drug and diseases genes which was previously applied to protein interaction networks and known drug targets (5) **(Supplementary methods)**. Here, we used tissue-specific BNs and tested the mean shortest distance between drug DEGs and disease genes.

The performance of PharmOmics drug repositioning was assessed using hyperlipidemia as the first test case, as multiple known drugs are available as positive controls. Since hyperlipidemia is most relevant to LDL and liver tissue, we retrieved LDL causal genes and pathways in liver tissue based on LDL GWAS and liver genetic regulation of gene expression using Mergeomics (**Supplementary methods**) (38), a method that can extract causal genes, pathways, and networks for diseases (39,40). In addition to retrieving disease genes from GWAS, a hyperlipidemia signature from CTD (16) was also used as an alternative source. For each drug with different dose and treatment durations, the signature with the highest overlap with the disease signature was used to represent the drug. Gene overlap- and network-based methods using dose/time-segregated signatures had similar overall performance (~90% AUC) in identification of antihyperlipidemic drugs (**Figure 5A, 5B)**, and the dose/time-segregated signatures performed better than the meta signatures when using network-based repositioning (**Figure 5C-5D)**. When compared to other platforms, PharmOmics was able to retrieve higher prediction rankings for the known drugs (**Table 1**) and better AUC (**Figure 5C-5D)** than CMAP and L1000 and higher balanced accuracy (**Supplementary Table 3**) than CREEDS, CMAP, and L1000. These results support the capacity of the PharmOmics platform as a drug repositioning tool.

**Table 1.**
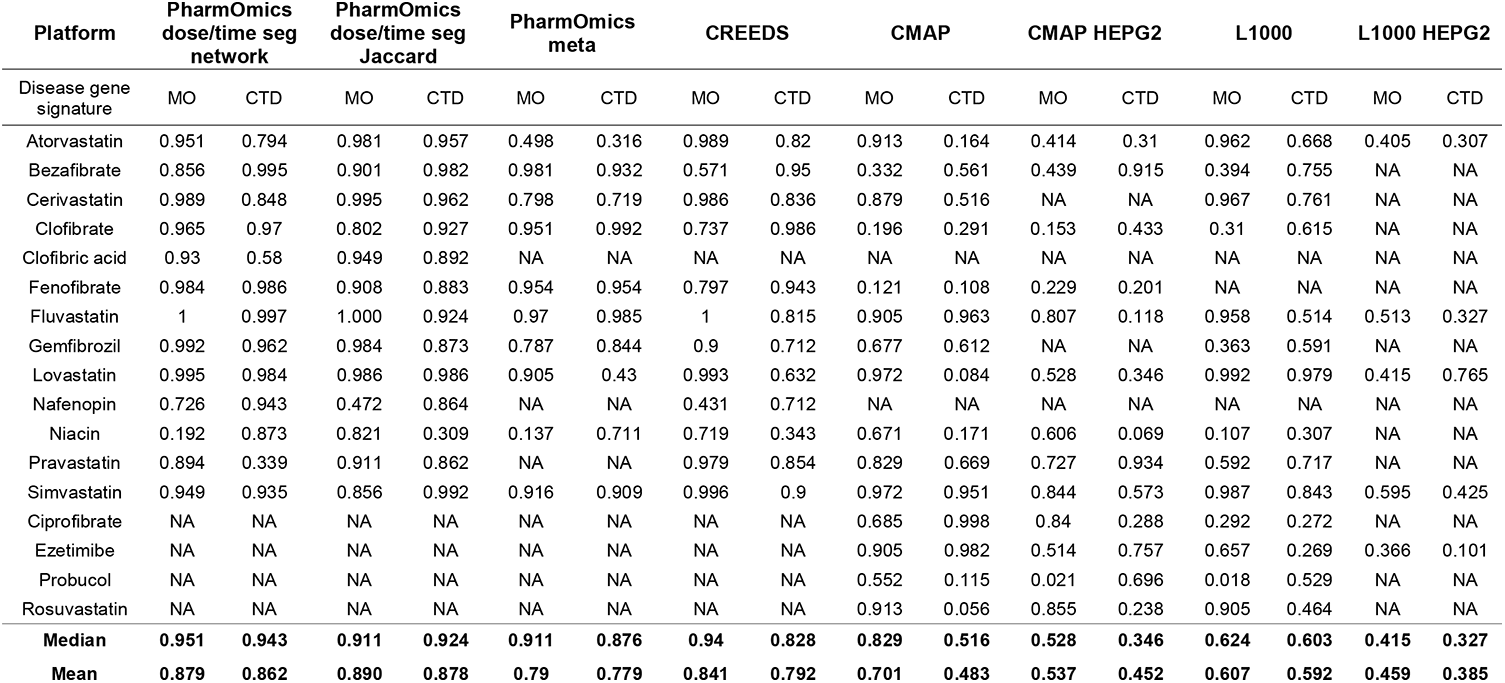
Prediction percentile of FDA approved antihyperlipidemic drug based on hyperlipidemia signatures from MergeOmics (MO) pipeline and atabase across different platforms tested. HEPG2 results from both L1000 and CMAP were retrieved for tissue specificity comparison.

| Platform | PharmOmics<br>dose/time seg<br>network |  | PharmOmics<br>dose/time seg<br>Jaccard |  | PharmOmics<br>meta |  | CREEDS |  | CMAP |  | CMAP HEPG2 |  | L1000 |  | L1000 HEPG2 |  |
| --- | --- | --- | --- | --- | --- | --- | --- | --- | --- | --- | --- | --- | --- | --- | --- | --- |
| Disease gene<br>signature | MO | CTD | MO | CTD | MO | CTD | MO | CTD | MO | CTD | MO | CTD | MO | CTD | MO | CTD |
| Atorvastatin | 0.951 | 0.794 | 0.981 | 0.957 | 0.498 | 0.316 | 0.989 | 0.82 | 0.913 | 0.164 | 0.414 | 0.31 | 0.962 | 0.668 | 0.405 | 0.307 |
| Bezafibrate | 0.856 | 0.995 | 0.901 | 0.982 | 0.981 | 0.932 | 0.571 | 0.95 | 0.332 | 0.561 | 0.439 | 0.915 | 0.394 | 0.755 | NA | NA |
| Cerivastatin | 0.989 | 0.848 | 0.995 | 0.962 | 0.798 | 0.719 | 0.986 | 0.836 | 0.879 | 0.516 | NA | NA | 0.967 | 0.761 | NA | NA |
| Clofibrate | 0.965 | 0.97 | 0.802 | 0.927 | 0.951 | 0.992 | 0.737 | 0.986 | 0.196 | 0.291 | 0.153 | 0.433 | 0.31 | 0.615 | NA | NA |
| Clofibric acid | 0.93 | 0.58 | 0.949 | 0.892 | NA | NA | NA | NA | NA | NA | NA | NA | NA | NA | NA | NA |
| Fenofibrate | 0.984 | 0.986 | 0.908 | 0.883 | 0.954 | 0.954 | 0.797 | 0.943 | 0.121 | 0.108 | 0.229 | 0.201 | NA | NA | NA | NA |
| Fluvastatin | 1 | 0.997 | 1.000 | 0.924 | 0.97 | 0.985 | 1 | 0.815 | 0.905 | 0.963 | 0.807 | 0.118 | 0.958 | 0.514 | 0.513 | 0.327 |
| Gemfibrozil | 0.992 | 0.962 | 0.984 | 0.873 | 0.787 | 0.844 | 0.9 | 0.712 | 0.677 | 0.612 | NA | NA | 0.363 | 0.591 | NA | NA |
| Lovastatin | 0.995 | 0.984 | 0.986 | 0.986 | 0.905 | 0.43 | 0.993 | 0.632 | 0.972 | 0.084 | 0.528 | 0.346 | 0.992 | 0.979 | 0.415 | 0.765 |
| Nafenopin | 0.726 | 0.943 | 0.472 | 0.864 | NA | NA | 0.431 | 0.712 | NA | NA | NA | NA | NA | NA | NA | NA |
| Niacin | 0.192 | 0.873 | 0.821 | 0.309 | 0.137 | 0.711 | 0.719 | 0.343 | 0.671 | 0.171 | 0.606 | 0.069 | 0.107 | 0.307 | NA | NA |
| Pravastatin | 0.894 | 0.339 | 0.911 | 0.862 | NA | NA | 0.979 | 0.854 | 0.829 | 0.669 | 0.727 | 0.934 | 0.592 | 0.717 | NA | NA |
| Simvastatin | 0.949 | 0.935 | 0.856 | 0.992 | 0.916 | 0.909 | 0.996 | 0.9 | 0.972 | 0.951 | 0.844 | 0.573 | 0.987 | 0.843 | 0.595 | 0.425 |
| Ciprofibrate | NA | NA | NA | NA | NA | NA | NA | NA | 0.685 | 0.998 | 0.84 | 0.288 | 0.292 | 0.272 | NA | NA |
| Ezetimibe | NA | NA | NA | NA | NA | NA | NA | NA | 0.905 | 0.982 | 0.514 | 0.757 | 0.657 | 0.269 | 0.366 | 0.101 |
| Probucol | NA | NA | NA | NA | NA | NA | NA | NA | 0.552 | 0.115 | 0.021 | 0.696 | 0.018 | 0.529 | NA | NA |
| Rosuvastatin | NA | NA | NA | NA | NA | NA | NA | NA | 0.913 | 0.056 | 0.855 | 0.238 | 0.905 | 0.464 | NA | NA |
| **Median** | **0.951** | **0.943** | **0.911** | **0.924** | **0.911** | **0.876** | **0.94** | **0.828** | **0.829** | **0.516** | **0.528** | **0.346** | **0.624** | **0.603** | **0.415** | **0.327** |
| **Mean** | **0.879** | **0.862** | **0.890** | **0.878** | **0.79** | **0.779** | **0.841** | **0.792** | **0.701** | **0.483** | **0.537** | **0.452** | **0.607** | **0.592** | **0.459** | **0.385** |

**Figure 5.**
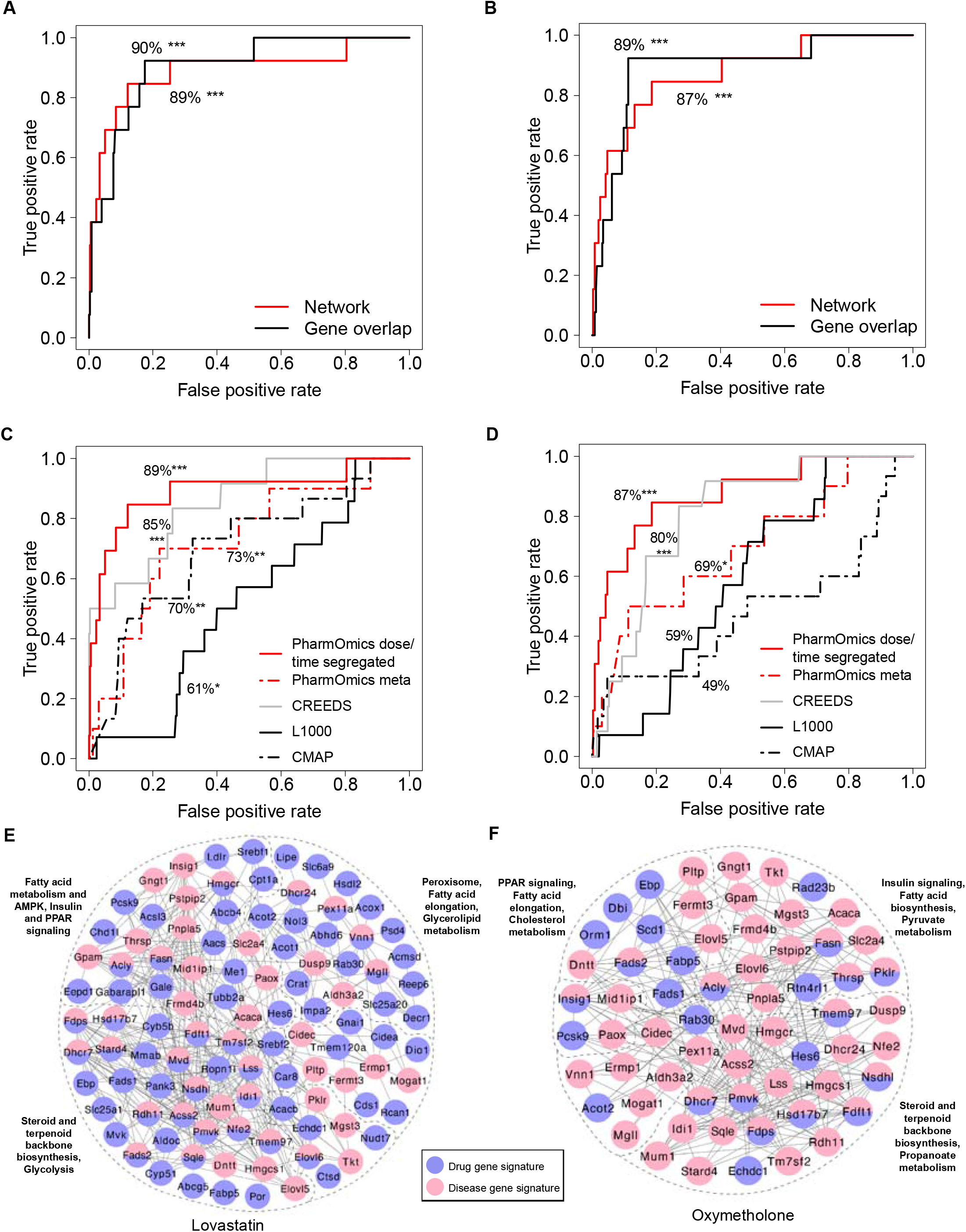
Drug repositioning for hyperlipidemia. AUC plots for network-based repositioning and gene overlap-based repositioning in identifying anti-hyperlipidemia drugs against other drugs using (A) Mergeomics hyperlipidemia signature or (B) CTD hyperlipidemia signature. Comparison of drug repositioning performance between PharmOmics network-based approach with CREEDS (using the “combined score” generated by the enrichment analysis tool implemented in Enrichr), L1000, and CMAP query system using (C) Mergeomics hyperlipidemia signature and (D) CTD hyperlipidemia signature. For drugs with multiple datasets with different doses and treatment times, only the best performing signature was used. (E) Drug-disease subnetwork of Mergeomics hyperlipidemia signature (red) and lovastatin signature (blue) showing first neighbor (direct) connections. (F) Drug-disease subnetwork Mergeomics hyperlipidemia signature (red) and oxymetholone signature (blue) showing first neighbor connections. Wilcoxon signed rank test was used to calculate significance between gene overlap/network z-scores between groups. *, **, *** indicates p < 0.05, p < 0.01 and p < 0.001 repectively.

We also examined the network overlap patterns of the top drugs consistently retrieved by the PharmOmics platform, lovastatin (ranked top 1% in both PharmOmics and CREEDS) and oxymetholone (ranked top 1% in PharmOmics and ranked as 15% in CREEDS). Both drugs targeted lipid metabolism genes (e.g. *Sqle* and *Hmgcr*) and PPAR pathways in the hyperlipidemia network (**Figure 5E, 5F**), but more lovastatin DEGs connected to disease genes compared to oxymetholone DEGs. These results support the utility of a network-based drug repositioning approach that does not require the direct retrieval of a known drug target or direct overlap of drug DEGs with disease genes.

We further evaluated the performance of PharmOmics in retrieving known drugs for other diseases. Using CTD disease signatures for hyperuricemia, we found network-based repositioning obtained 90% AUC (p=0.009) for detection of anti-hyperuricemia drugs, whereas the gene overlap-based method did not yield a significant AUC (prediction ranks in **Supplementary Table 4**). We also queried hepatitis signatures and achieved 83% AUC (p< 0.001) using the network method and 79% AUC (p<0.001) using the gene overlap method in retrieving non-steroid anti-inflammatory agents (prediction ranks in **Supplementary Table 5)**. Finally, using diabetes signatures, PharmOmics was able to predict PPAR gamma agonist drugs (79% AUC, p=0.04), but not sulfonylurea drugs which act on the pancreatic islet to enhance insulin release (prediction ranks in **Supplementary Table 6**). We note that the paucity of drug signatures in diabetes relevant tissues/cells such as the islets and the digestive system likely explains why sulfonylurea drugs are harder to retrieve. Overall, these various test cases using known therapeutic drugs as positive controls support the utility of network-based drug repositioning for select diseases when drug signatures from the appropriate tissues are used.

### Use of PharmOmics to predict drugs for NAFLD

After establishing the performance of PharmOmics in drug repositioning using the case studies above, we applied PharmOmics to predict potential drugs for NAFLD, for which there is currently no approved drugs. Using NAFLD steatosis signatures from a published study (40) and the CTD database (16), we predicted PPAR alpha agonists (clofibrate, fenofibrate, bezafibrate, and gemfibrozil), HMG-CoA reductase inhibitors (lovastatin, fluvastatin, and simvastatin), and a PPAR gamma agonist (rosiglitazone) among the top 10% of the drug candidates (**Supplementary Table 7**). PPAR agonists have been supported as potential drugs for NAFLD (41–52). Statins have shown efficacy in animal models (34,53), although clinical results are controversial (54,55). Additional predicted drugs included aspirin, which was recently reported to be associated with reducing liver fibrosis progression (56).

### In vivo validation of drug repositioning predictions for NAFLD

Next, we sought to experimentally validate the ability of two top ranked drugs by PharmOmics, fluvastatin and aspirin, to mitigate liver steatosis as predicted by PharmOmics and assess the accuracy of repositioning ranks. Compared to other platforms (**Supplementary Table 7**), fluvastatin was ranked high consistently in PharmOmics (top 5%), CMAP (top 1% in all cells combined, 20% in HEPG2), CREEDS (20%), and L1000 (top 1% in all cells combined, 55% in HEPG2). In comparison, aspirin was ranked higher in PharmOmics (top 5%) compared to CREEDS (30%) and CMAP (35%) and was not documented in L1000. Therefore, these predictions are relatively unique to PharmOmics.

Comparison between the mice in HFHS group (NAFLD) and the chow group (Control) confirmed HFHS induced NAFLD phenotypes including increased body weight, adiposity, and hepatic steatosis (**Supplementary Figure 2A and 2B**). Comparison of the fluvastatin and aspirin treated groups with the NAFLD group revealed significant drug effects on body weight gain for both fluvastatin (p<0.0001; **Figure 6A**) and aspirin (p<0.0001; **Figure 6B**). The adiposity phenotype (fat and lean mass ratio) also showed significant drug effects from both fluvastatin (p<0.0001; **Figure 6C**) and aspirin (p=0.0157; **Figure 6D**).

**Figure 6.**
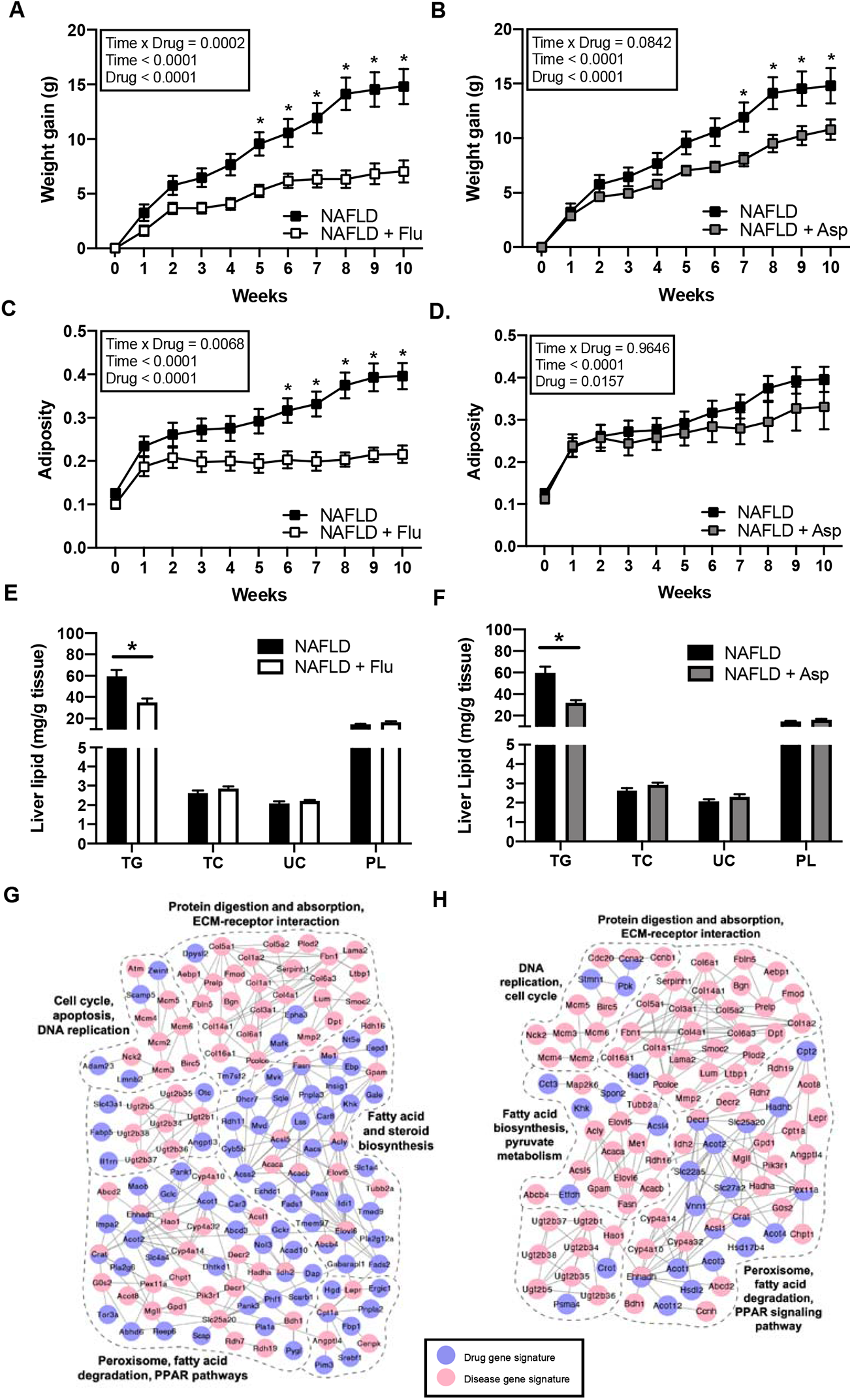
*In vivo* validation of predicted drugs Fluvastatin and Aspirin on preventing NAFLD phenotypes in C57BL/6J mice. (A and B) Time course of body weight gain in mice treated with fluvastatin (A) and aspirin (B) over 10 weeks. (C and D) Time course of fat mass and muscle mass ratio (adiposity) in mice treated with fluvastatin (C) and aspirin (D) over 10 weeks. (A-D) Data were analyzed by two-way ANOVA followed by Sidak post-hoc analysis to examine treatment effects at individual time points. P value < 0.05 was considered significant and is denoted by an asterisk (*). (E and F) Quantification of lipids in the liver of mice on fluvastatin (E) and aspirin (F) treatment for 10 weeks. Triglyceride (TG), Total Cholesterol (TC), Unesterified Cholesterol (UC), Phospholipid (PL). (D and E) Data were analyzed using two-sided Student’s t-test. P value < 0.05 was considered significant and is denoted by an asterisk (*). Sample size nLJ=LJ7-9/group. High fat high sucrose (HFHS) group (NAFLD); HFHS with fluvastatin (NAFLD + Flu); HFHS with aspirin (NAFLD + Asp). (G-H) Gene network view of fluvastatin gene signatures overlapping with NAFLD disease signatures (G) Gene network view of aspirin gene signatures overlapping with NAFLD disease signatures (H).

There was no significant difference in total liver weight among the groups (**Supplementary Figure 2C** for Control and NAFLD group comparison; **Supplementary Figure 3A and 3B** for NAFLD and drug group comparisons). As expected, the HFHS group had significantly elevated levels of liver TG compared to controls, without changes in other lipids measured such as TC, UC, and PL (**Supplementary Figure 2D**). In the drug treatment groups, both fluvastatin (p=0.0044) and aspirin (p=0.0023) induced significant decreases in hepatic TG compared to the NAFLD group, without any effect on TC, UC, and PL (**Figure 5E-5F**).

We further investigated whether the effects of the drugs on NAFLD phenotypes were confounded by food and water intake. No effect of food intake was observed in the NAFLD + Flu group; however, there was a significant decrease in food intake in the NAFLD + Asp group (**Supplementary Figure 3C-3D**). No effect on water intake was found for both groups (**Supplementary Figure 3E-3F**). We next adjusted for food intake in the NAFLD phenotypic analysis for body weight gain, adiposity, and TG levels using linear regression. After the adjustment, the significant effects of fluvastatin on NAFLD phenotypes remained (body weight gain p=0.0306; adiposity p=0.0022; hepatic TG p=0.0190). For aspirin, the significant effects on adiposity (p=0.0479) and hepatic TG (p=0.0372) remained, but the effect on body weight gain was no longer significant (p=0.0559). Overall, food/water intake did not have major influence on treatment effects on NAFLD observed for both drugs.

Our experimental validation experiments support the efficacy of both fluvastatin and aspirin in mitigating NAFLD. The effects of fluvastatin were stronger than that of aspirin and visualization of the network overlaps between NAFLD signatures and drug signatures revealed more extensive disease network connections for fluvastatin than for aspirin (**Figure 6G-6H**), supporting their repositioning ranks and potential mechanisms of action. The signatures of the two drugs connected to pathways involved in NAFLD such as PPAR signaling pathways, fatty acid and steroid biosynthesis (**Figure 6G-6H**).

### Utility of PharmOmics drug signatures in predicting and understanding hepatotoxicity

We further explored the potential of coupling PharmOmics drug signatures and tissue networks to predict liver toxicity, a major type of ADR for which both toxicity signatures and orthogonal ADR documentations from various independent databases are available for performance evaluation. We used the chemical-induced liver injury signature containing 435 genes from CTD to match with PharmOmics drug signatures through liver gene networks. We then used both the histological severity from TG-GATEs and the independent FDA drug-induced liver disease (DILI) categories (“most”, “less” – moderate/mild DILI adverse reactions compared to the “most” category, and “no” DILI concern) as *in silico* independent validation of the drugs predicted by PharmOmics that match with the CTD liver toxicity signature.

First, we examined the relationship between the matching scores of PharmOmics signatures and the histological severity grading based on TG-GATEs. Both the network-based and gene overlap-based scores from PharmOmics increased with higher histological severity defined by TG-GATEs (**Figure 7A**). Next, we examined the dose-dependent effects across the TG-GATEs histological severity categories as well as the three FDA DILI categories. Our results indicated that severe histological grading occurred mainly at higher drug doses within both the “less” and “most” DILI concern categories (**Figure 7B**). Analysis of the relationship between dose/time-segregated signatures and network-based PharmOmics scores indicated that drug treatment at higher doses had higher network matching ranks in PharmOmics and more severe DILI (**Figure 7C**). In addition, we tested the performance of PharmOmics in predicting hepatotoxic drugs from the FDA DILI drug database. PharmOmics dose/time-segregated signatures resulted in higher performance (67% AUC) compared to the meta signatures (60% AUC) and the other platforms tested such as CREEDS, CMAP, and L1000 (AUC 50-53%; **Figure 7D; Supplementary Figure 4**). Top drug predictions based on the complete hepatotoxicity signatures were wy-14643 (experimental drug with severe histological finding in TG-GATEs), dexamethasone (moderate DILI concern category in FDA and moderate histological finding in TG-GATEs), phenobarbital (moderate DILI concern), indomethacin (“most” DILI concern), and fenofibrate (moderate DILI concern).

**Figure 7.**
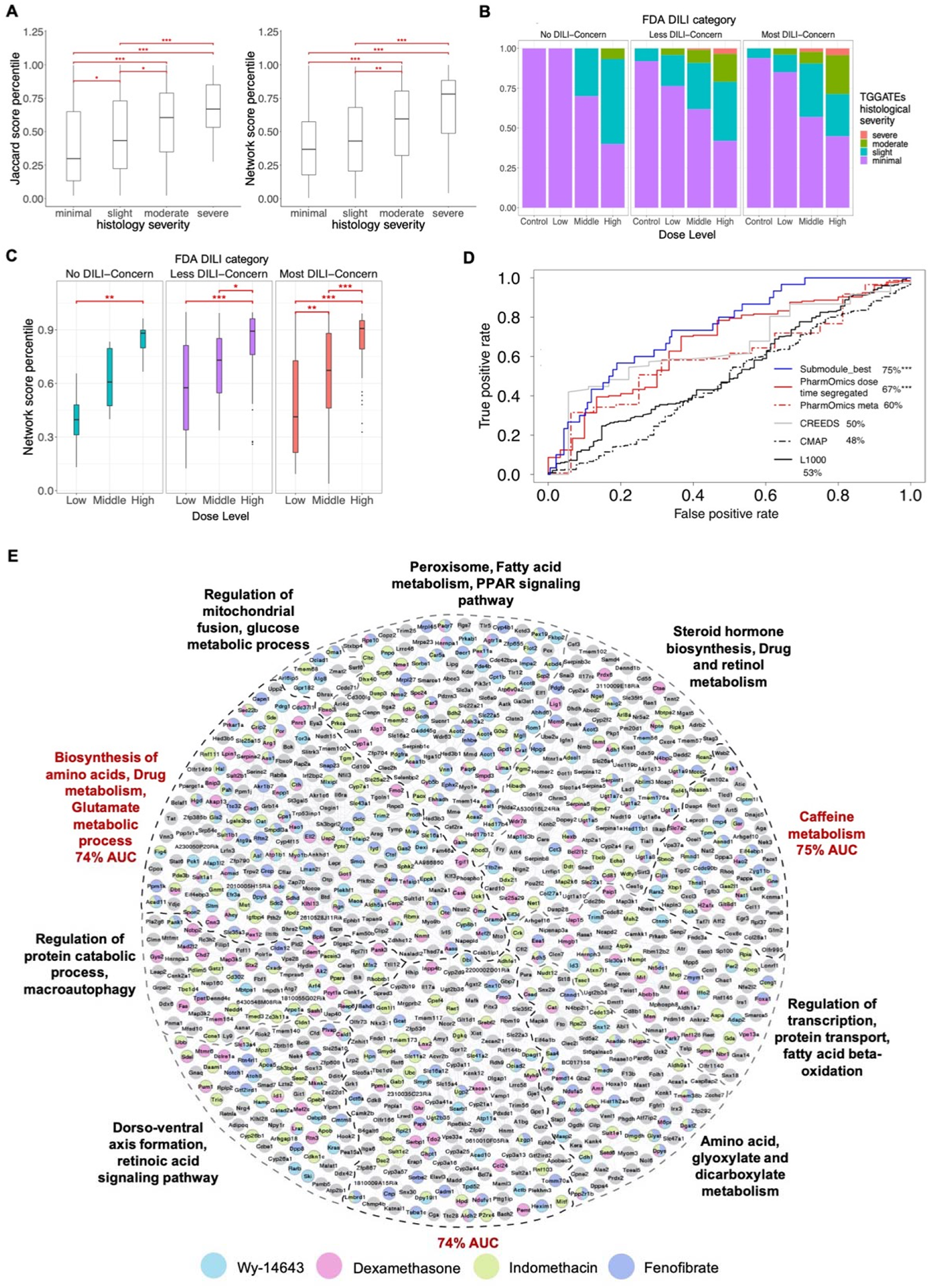
Utility of PharmOmics drug signatures in hepatotoxicity prediction based on matching between PharmOmics drug signatures and hepatotoxicity signatures of drug induced liver injury (DILI) curated from comparative toxicogenomics database (CTD). (A) Boxplots of Jaccard score-based hepatotoxicity ranking (left) and network-based hepatoxicity ranking (right) by PharmOmics, across four categories of liver injury histological severity defined by the independent TG-GATEs database (x-axis). PharmOmics hepatotoxicity scores are higher for more severe liver injury categories. (B) PharmOmics hepatotoxicity prediction scores based on gene signatures of higher drug doses correspond to more severe liver injury categories defined by TG-GATES across three DILI concern categories (“no”, “less”, “most”) defined by FDA. (C) Boxplots of network-based hepatoxicity scores show increased scores at higher doses across three FDA DILI concern categories. (D) ROC curves comparing PharmOmics with other tools in predicting hepatotoxic drugs from the FDA DILI drug database. For PharmOmics, three sets of tests were performed, where dose/time-segregated drug signatures, meta signatures, or a hepatotoxicity subnetwork was used. (E) Liver hepatotoxicity network based on CTD hepatotoxicity genes and its overlap with drug signatures of 4 of the top 5 predicted drugs by PharmOmics which had >50 signature genes. Phenobarbital was among the top 5 drugs but was not included in the figure due to its small DEG size. Colors of the network nodes denote the different drugs targeting the genes. The top 3 predictive subnetworks are depicted in red (D). ANOVA test followed by post-hoc analysis was used for statistics in A and C. *, **, *** indicates p < 0.05, p < 0.01 and p < 0.001 respectively. Boxplots show interquartile range (IQR) and median values (line inside the box). IQR was defined as between 25th (Q1) and 75th (Q3) percentile. The upper and lower bars indicate the points within Q3 + 1.5*IQR and Q1 – 1.5*IQR, respectively.

Since CTD curated a large number of genes (435 genes) related to chemical induced liver injury, we hypothesized that this large network could be divided into subnetworks indicative of different mechanisms towards liver toxicity, which could improve toxicity prediction for drugs with different mechanisms. We first examined network overlapping patterns of the top 5 predicted drugs by using the CTD liver injury genes (**Figure 7E**) and found consistent targeting of gene subnetworks across top predictions. We then applied the Louvain clustering method to divide the liver injury network into subnetworks and filtered subnetworks with less than 10 genes to reduce uncertainty. These different subnetworks showed varying abilities in identifying drugs with DILI concerns (**Supplementary Table 8**). The best performing subnetwork showed improved AUC compared to the whole network (75% vs 67%; **Figure 7D**). Further scrutinization of key genes documented in CTD signatures of the top performing subnetwork revealed that the antioxidant gene *GSR*, the phase 2 drug metabolizer *NAT2*, and the inflammatory response gene *IRAK1* showed the best predictability (**Supplementary Table 8**). These results suggest that the network-based toxicity prediction approach may help prioritize predictive genes, pathways, and subnetworks related to hepatotoxicity.

### Utility of PharmOmics drug signatures in predicting and understanding nephrotoxicity

We also examined the performance of PharmOmics in predicting nephrotoxicity, another ADR for which both toxicity signature and drug ADR documentations are available from independent sources to help validate performance.

Nephrotoxicity signatures were curated from CTD using either the chronic kidney disease signature (CKD, 56 genes) or acute kidney injury signature (AKI, 120 genes), which were matched with PharmOmics drug signatures to predict drugs matching CKD or AKI signatures. The PharmOmics predictions were then validated using kidney histological severity documented by TG-GATEs or nephrotoxicity defined by DrugBank. There were 13 shared genes between CKD and AKI signatures including several inflammatory factors *TNF, TGFB1, NFKB1,* and *IL6*. We found that unlike AKI signatures, using CKD signatures against PharmOmics drug signatures resulted in network matching scores (not gene overlap scores) that agreed well with histological severity documented in TG-GATEs (**Figure 8A**). Therefore, we focused on using CKD signatures in downstream network-based analyses. We found that PharmOmics drug signatures of higher doses predicted more drugs with severe or moderate kidney histology categorized by TG-GATEs as well as drugs with nephrotoxicity as defined by DrugBank (**Figure 8B**). However, when examining the relationship between PharmOmics network scores across doses and DrugBank nephrotoxicity categories (non-nephrotoxic or nephrotoxic), the network scores did not show a significant dose-dependent relationship (**Figure 8C**). This is in contrast to the dose-dependent relationship observed for hepatotoxicity analysis (**Figure 7C**). The weaker performance of PharmOmics in nephrotoxicity prediction could be due to the smaller number of kidney drug datasets (~1k) compared to liver drug datasets (~5k) based on data availability.

**Figure 8.**
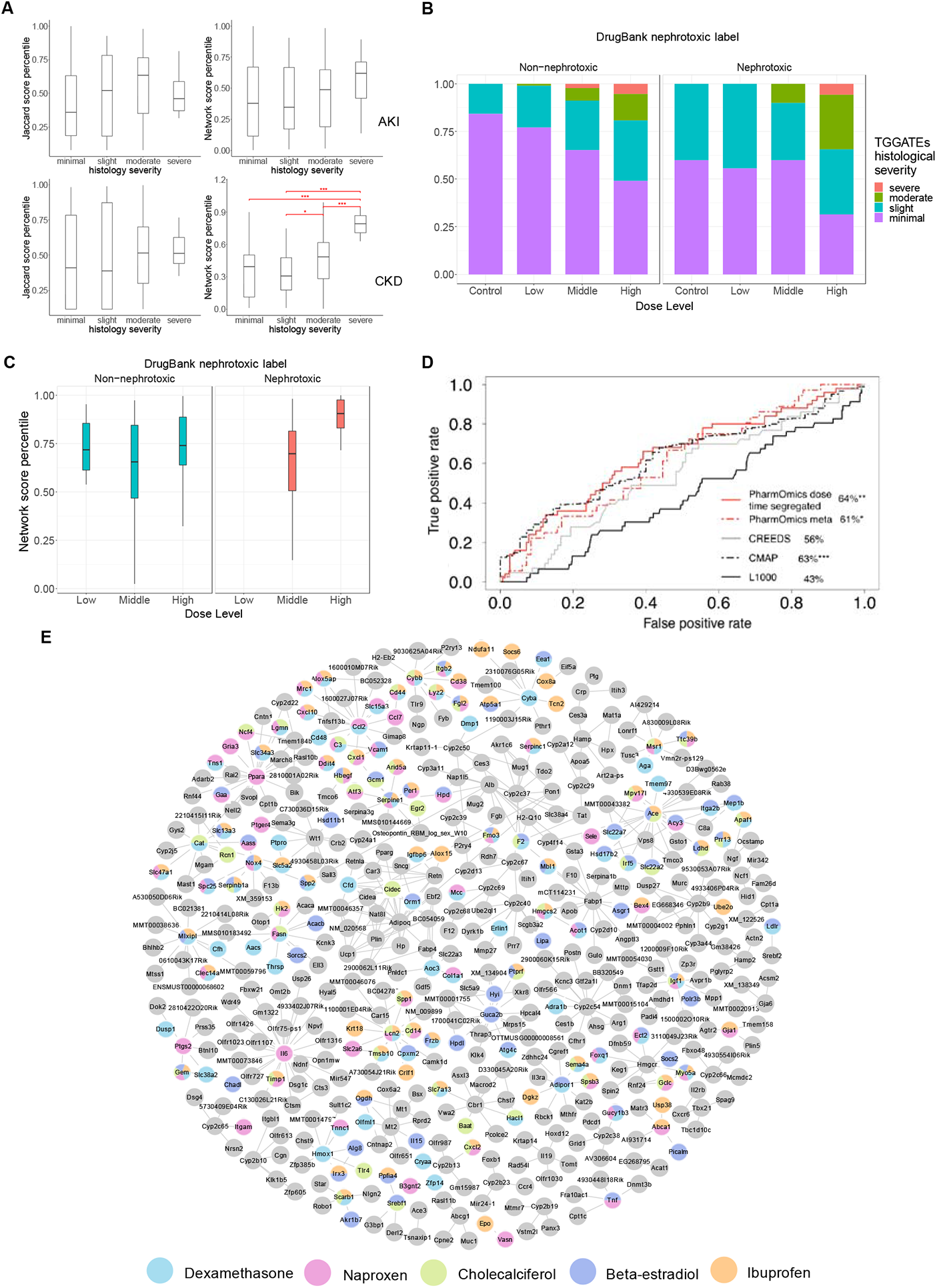
Utility of PharmOmics drug signatures in nephrotoxicity prediction based on matching between PharmOmics drug signatures and nephrotoxicity signatures of chemical induced acute kidney injury (AKI) or chronic kidney disease (CKD) from comparative toxicogenomics database (CTD). (A) Boxplots of Jaccard score-based nephrotoxicity ranking (left) and network-based nephrotoxicity ranking (right) by PharmOmics, based on matching with AKI (top) or CKD (bottom) nephrotoxicity genes from CTD, across four categories of kidney histological severity defined by the independent TG-GATEs database. Network-based nephrotoxicity prediction by PharmOmics showed a positive relationship between nephrotoxicity scores by PharmOmics and kidney histology severity defined by TG-GATEs. (B) PharmOmics nephrotoxicity prediction scores based on gene signatures of higher drug doses correspond to more severe kidney injury categories defined by TG-GATES, segregated by nephrotoxic labels defined by DrugBank. (C) Boxplot of network-based nephrotoxicity scores, using CKD nephrotoxicity genes against PharmOmics drug signatures, did not show significant dose-dependent trend in non-nephrotoxic drugs or nephrotoxic drugs defined by DrugBank. The low dose treatment group for the nephrotoxic drugs did not contain drug signatures with more than 10 genes and therefore the scores were not plotted. (D) ROC curve comparing the performance of PharmOmics with other tools in predicting nephrotoxic agents in DrugBank. For PharmOmics, two sets of tests were performed, where either dose/time-segregated drug signatures or meta signatures was used. (E) Kidney nephrotoxicity network based on CTD nephrotoxicity genes and the network overlap with drug signatures of top 5 drugs predicted by PharmOmics. Colors of the network nodes denote the various drugs targeting the genes. ANOVA test with post-hoc analysis was used for statistics in A and C. *, **, *** indicates p < 0.05, p < 0.01 and p < 0.001 respectively. Boxplots show interquartile range (IQR) and median values (line inside the box), with IQR defined as between 25th (Q1) and 75th (Q3) percentile.

Finally, we assessed the performance of PharmOmics and other tools in identifying DrugBank nephrotoxic agents. PharmOmics dose/time-segregated (64% AUC, p=0.001) and meta databases (61% AUC, p=0.028) both showed a significant performance (**Figure 8D**), whereas from the other tools evaluated only CMAP (63% AUC, p<0.001) showed a significant performance (L1000 43% and CREEDS 56% AUC, non-significant). The top 5 nephrotoxic drugs predicted by PharmOmics were dexamethasone (potential CKD alleviating agent) (57,58), naproxen (documented nephrotoxic drug in DrugBank), cholecalciferol (potential CKD alleviating agent) (59), beta-estradiol (potential alleviating agent in women) (60,61), and ibuprofen (documented nephrotoxic drug in DrugBank). We also examined the gene overlap patterns of the top drugs with the CKD gene network (**Figure 8E**) and found sparse overlap, which again is in contrast to the top hepatotoxicity drugs (**Figure 7E**).

Overall, our assessment of the application of PharmOmics in toxicity or ADR prediction supports its potential value but also emphasizes that PharmOmics drug signatures may have differing performance in different use cases. Several factors, including the toxicity signatures used (e.g., CKD signature performed better than AKI signature for nephrotoxicity prediction), ADR/toxicity annotation (e.g., TG-GATEs or DrugBank), and signature matching method (network-based approach better than gene overlap approach) can all significantly affect the results. We also note that our network approach does not differentiate toxicity-inducing drugs from toxicity-mitigating drugs since it is based on network connectivity and not the directionality of gene signatures.

### Utility of meta-analysis signatures to understand tissue and species specificity

We used meta signatures, which reflect the dose-independent, consistent genes affected by drugs across studies in the same tissue or species, to evaluate tissue and species specificity of drugs by analyzing the overlap in gene signatures for each drug across different tissues and species and visualized the results using UpSetR (62). As shown in **Figure 9A**, the overlap rate in the DEGs of the same drug between tissues and organs is usually less than 5%, indicating a high variability in DEGs between tissues. As an example, we examined atorvastatin, a HMGCR (β-Hydroxy β-methylglutaryl-CoA receptor) inhibitor, which has well understood mechanisms and has been broadly tested in different tissues under the human species label. We found that two DEGs (*TSC22D3, THBS1*) were shared across tissues (**Figure 9B**). These genes are involved in extracellular matrix and inflammation, suggesting these processes are common targets of atorvastatin across tissues. Among the pathways shared across tissues, immune related pathways were shared between blood cells and liver cells but not in prostate cells from the urogenital system (**Figure 9C, Supplementary Table 9)**. Pathway analysis indicated that steroid synthesis and drug metabolism pathways were altered primarily in liver, which is expected as the known target of statin drugs is HMGCR, the rate limiting enzyme in cholesterol biosynthesis in liver. Blood monocyte DEGs indicated changes in inflammation related pathways, while GPCR ligand binding proteins were altered in prostate cancer cells. The tissue specificity of drug meta signatures revealed through our analysis supports tissue-specific therapeutic responses and side effects and emphasizes the need for comprehensive inclusion of drug signatures from different tissue systems as implemented in the PharmOmics framework.

**Figure 9.**
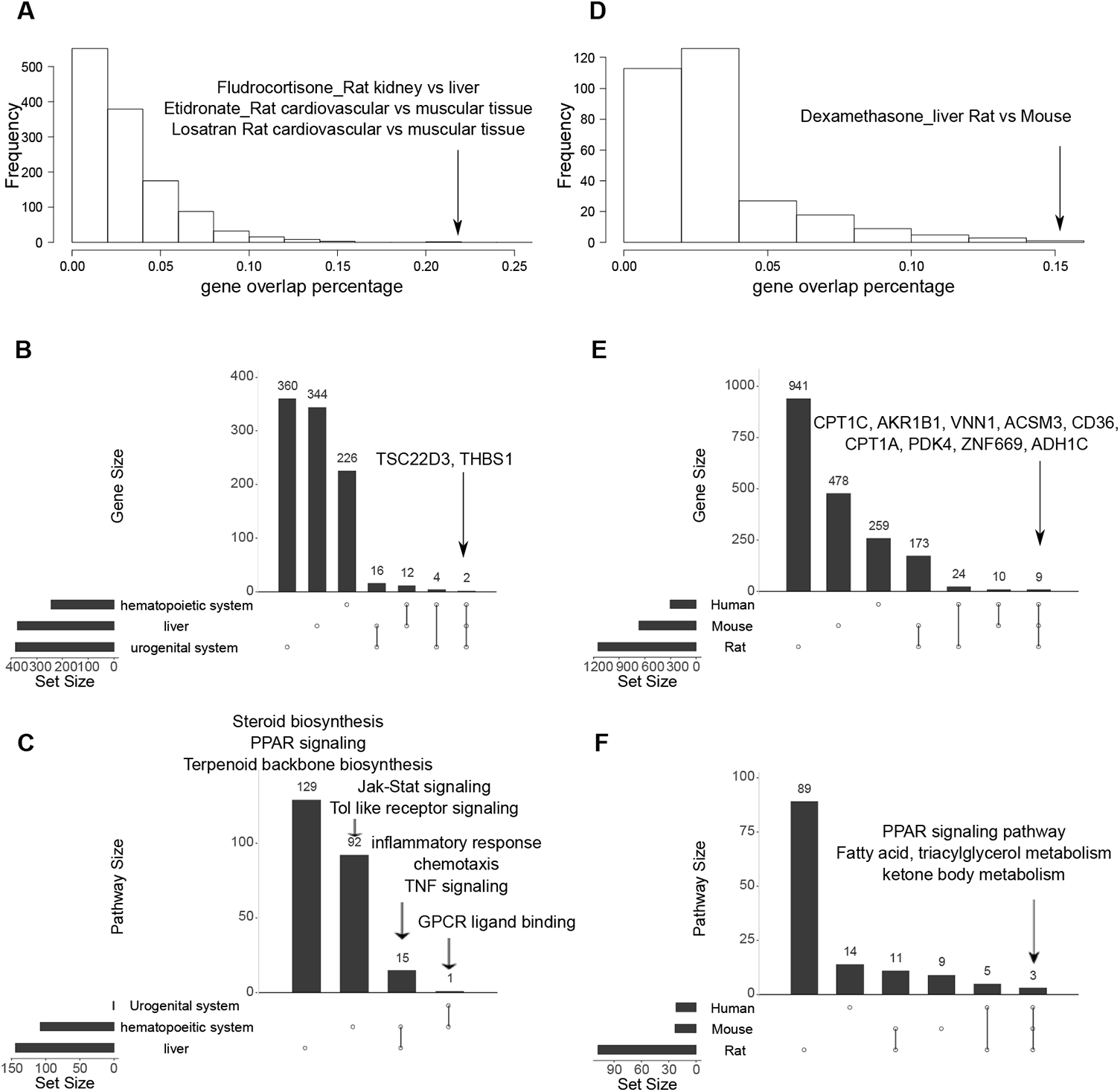
Cross-tissue and cross-species comparisons of drug signatures in PharmOmics. (A) Distribution of drug signature overlap percentages between tissue pairs in matching species from PharmOmics. Arrow points to the pairs of tissues for drugs with high overlap in gene signatures. (B) Upset plot of cross-tissue comparison for atorvastatin signatures genes. Y-axis indicates number of genes. (C) Upset plot of cross-tissue comparison for pathways enriched in atorvastatin signatures. Y-axis indicates number of pathways. (D) Distribution of drug signature overlap percentages between pairs of species for matching tissues from PharmOmics. Arrow points to the species pair with high gene signature overlap for a matching drug. (E) Upset plot of cross-species comparison for rosiglitazone liver gene signatures. (F) Upset plot of cross-species comparison for pathways enriched in rosiglitazone liver signatures. Pairs of tissues with shared drug signature genes or pathways are connected with black vertical lines in the bottom portion of the Upset plots.

We also found evidence for high species specificity. As shown in **Figure 9D**, the pair-wise overlaps in DEGs between species for the same drug is generally lower than 5%. Here we chose PPAR gamma receptor agonist rosiglitazone as an example because this drug has datasets across human, rat, and mouse in PharmOmics, and its mode of actions is well-studied. As shown in **Figure 9E** and **9F**, nine genes (*CPT1C, AKR1B1, VNN1, ACSM3, CD36, CPT1A, PDK4, ZNF669, ADH1C*) and several pathways (PPAR signaling and fatty acid, triacylglycerol, and ketone body metabolism) were consistently identified from liver DEGs across species (**Supplementary Table 10**), reflecting the major species-independent pharmacological effects of rosiglitazone. Bile acid related genes were altered in rat datasets, whereas retinol metabolism and adipocytokine pathways were altered in human datasets. The species-specific differences identified highlight the importance of understanding the physiological differences among model systems to facilitate drug design with better translational potential. Our cross-species comparative studies also emphasize the need to investigate drugs in multiple species, as only 21% of the unique drug-tissue pairs (236 out of 1144) from PharmOmics meta signatures have data from two or more species.

## Discussion

Here we present PharmOmics, a publicly available drug signature database along with an open-access web interface for accessing and utilizing the signatures for various applications. PharmOmics utilizes published drug-related transcriptomic datasets across multiple data repositories and provides unique tissue-, species-, and dose/time-stratified gene signatures that are more reflective of *in vivo* activities of drugs. We also developed a unique framework for drug repositioning based on tissue-specific gene network models. We examined the potential applications of PharmOmics for various utilities including drug repurposing, toxicity prediction, target identification, and comparisons of molecular activities between tissues and species. We also carried out *in silico* performance assessments across drug signature databases and *in vivo* mouse experiments to validate our network-based drug predictions for NAFLD.

Compared to the well-established CMAP and LINC1000 platforms, PharmOmics focuses more on *in vivo* settings and likely captures more physiologically relevant drug signatures to improve drug repositioning performance. Compared to a previous crowdsourcing effort which also utilizes publicly available drug datasets (18), our PharmOmics platform included more curated databases (TGGATEs + drugMatrix Affymetrix + drugMatrix Codelink datasets compared to only drugMatrix Codelink datasets from CREEDS) and involved systematic tissue, species, and treatment regimen stratification to facilitate drug repositioning. Our platform is also the only tool utilizing a gene network framework rather than direct gene overlap approach.

The use of tissue annotation with Brenda Tissue Ontology helps normalize organ labels and improves comparability of datasets. The unique tissue- and species-specific analyses implemented in PharmOmics allows for comprehensive molecular insight into the actions of drug molecules in individual tissues and species. Our results support that different species have unique drug responses in addition to shared features; therefore, drug responses obtained in animal models require caution when translating to humans. This notion agrees with the long-observed high failure rate of drug development that has primarily relied on preclinical animal models and argues for greater consideration and understanding of inter-species differences in drug actions.

In addition to tissue and species stratification, we also provide detailed dose/time-segregated drug signatures, which can help better understand the dose- and time-dependent effects of drugs through gene signature and pathway comparisons offered through our web server. By contrast, the meta-analysis signatures capture the consistent genes and pathways across treatment regimens, which likely represent core, dose/time-independent mechanisms, and help address the sample size issue of individual datasets since the majority of drug treatment datasets have n<=3/group. Dose/time-segregated signatures performed better than meta signatures for both drug repositioning and toxicity prediction. However, meta signatures showed better performance than CMAP, LINC1000, and CREEDs **(Figure 5, 7, 8)**, and can also significantly shorten the computation time in network-based repositioning applications. For instance, computation using 1251 human meta signatures can be completed in 40 minutes, whereas using ~14,000 dose/time-segregated signatures can take 4 hours. These estimates will vary depending on input data size and server load.

Previous drug repositioning studies support the utility of a protein network-based approach for drug repositioning. Here we show that combining the drug transcriptomic signatures in PharmOmics with tissue-specific gene regulatory networks and gene signatures of diseases can retrieve known therapeutic drugs, predict potential therapeutic avenues, and predict tissue toxicity. Compared to other platforms, the use of tissue- and species-specific drug signatures along with network biology is a unique strength of PharmOmics, which enables drug prioritization based on network proximity rather than direct gene overlaps. We demonstrate in various applications that network-based analysis had a superior performance to that of gene overlap-based analysis. Moreover, the tissue-specific network connections between drugs and diseases or toxicity offer molecular and mechanistic insights into the therapeutic or toxic effects of drugs. For instance, fluvastatin showed different NAFLD overlapping patterns compared to aspirin, which inferred differences in disease repositioning depending on different drug mechanisms.

In general, gene signatures of drugs reflect cascades of downstream events after drug administration. The initial drug target(s) may or may not be captured by drug DEGs due to the lack of dynamic information in the DEGs. Therefore, we explored if PharmOmics signatures as well as signatures from other platforms can be used to retrieve drug targets through integration with pathway or network information. Our results show that DEGs may help inform on the pathways affected by the drugs but retrieving the direct targets can be difficult. We caution the use of drug DEGs from any drug signature platform for direct target identification.

There are several limitations in this study. First, our computational pipeline may not be able to identify all of the drug datasets from GEO and ArrayExpress database. Variations in annotations of drug names, sample size, definition of treatment vs control groups, and tissue/cell line labeling across datasets make it challenging to design a fully automated pipeline to curate drug signatures. It is therefore crucial for GEO and ArrayExpress repositories to offer clear definitions and instructions for metadata generation in order to standardize terms across datasets to facilitate future data reuse. Secondly, the coverage of tissue, species, and treatment regimens across drugs is unbalanced, preventing a thorough comparison across tissues, species, dosages, and treatment windows. We will continue to refine the pipeline and update our PharmOmics database periodically to include more datasets as they become available to increase the coverage of datasets and drug signatures. Thirdly, the sample sizes for drug treatment studies tend to be small (majority with n=3/group or less), which limit the statistical power and reliability of the drug signatures when individual studies were analyzed. This is an intrinsic limitation of existing drug studies and highlights the need for systematic efforts to increase sample sizes in such studies. To mitigate this concern and reduce the reliance on individual studies, we implemented a meta-analysis strategy to aggregate drug signatures from individual studies and derive meta signatures. However, this strategy removes dosage- and time-dependent effects. We offer both options in our database to mitigate sample size concerns through meta-analysis and retain dose and time regimen information through dose/time-segregated analysis. Fourth, our network-based applications are currently limited in the coverage of high-quality tissue specific regulatory networks and computational power. We will continue to expand and improve the tissue networks and computing environment in our web server. Lastly, systematic validation efforts are needed to substantiate the value of our platform. We utilized both *in silico* performance assessments and *in vivo* experiments to validate our predictions in limited settings. We mainly focused on liver related diseases with well-documented drugs and disease signatures (hyperlipidemia, hyperuricemia, diabetes, and liver/kidney toxicity) to benchmark the utilities of PharmOmics and experimentally validated two drugs predicted for NAFLD. As with the other existing platforms such as CMAP and LINC1000, future application studies and community-based validation efforts are necessary to assess the value of PharmOmics.

## Conclusion

We have established a new drug signature database, PharmOmics, across different species and tissues, which captures the systems level *in vivo* activities of drug molecules. In addition, we demonstrate the possible means to integrate these signatures with network biology to address drug repositioning needs for disease treatment and to predict and characterize liver and kidney injury. PharmOmics has the potential to complement other available drug signature databases to accelerate drug development and toxicology research. Our PharmOmics database and pipeline will be updated periodically to include newly available datasets to increase the coverage of the drug signatures across tissues and species. It should be noted that we aim to position PharmOmics as a data-driven compensatory tool in hypothesis generation. Integration with known drug characteristics to select drug candidates and design follow up experiments are still essential.

## List of abbreviations

ADR: adverse drug reactions
CTD: comparative toxicogenomics database
KEGG: Kyoto Encyclopedia of Genes and Genomes
DEG: differential expressed genes
FDR: false discovery rate
wKDA: weighted key driver analysis
NAFLD: non-alcoholic fatty liver disease
LDL: low-density lipoprotein cholesterol
GWAS: genome-wide association study
BN: Bayesian gene regulatory network
ROC: Receiver operating characteristic
HMGCR: β-Hydroxy β-methylglutaryl-CoA receptor
PPAR: Peroxisome proliferator-activated receptor
GPCR: G-protein coupled receptor

## Declaration

### Ethics approval and consent to participate

Not applicable.

### Consent for publication

Not applicable.

### Availability of data and materials

Indexed dataset catalog, pre-computed gene signatures and pre-computed pathway enrichments for individual drugs are deposited to and accessible through the PharmOmics web server (http://mergeomics.research.idre.ucla.edu/runpharmomics.php). We also implemented functions for same-tissue between-species comparison and same-species between-tissue comparison. Direct download of select drug signatures is also enabled. In addition, network-based drug repositioning analysis and gene overlap-based drug repositioning analysis using all drug signatures are available at http://mergeomics.research.idre.ucla.edu/runpharmomics.php.

### Authors’ contributions

YC curated and analyzed data, constructed database, and designed and conducted application studies. GD, JY, and PC conducted validation experiments. JD, TXN, DH, and MB designed and implemented the PharmOmics web server. DA provided support in data curation and analysis. GA, JG, NZ, and PP assisted with data curation. YC, GD, JD, and XY wrote the manuscript. XY designed and supervised the research. All authors contributed to manuscript editing.

## Acknowledgements

We thank Dr. Patrick Allard at UCLA for constructive input on the manuscript. We thank Dr. Avi Ma’ayan at Icahn School of Medicine at Mount Sinai for providing L1000FWD and CREEDS data for comparison.

