## Supplementary information, Supplementary Table 1-6,11 for "PharmOmics: A Species- and Tissue-specific Drug Signature Database and Online Tool for Drug Repurposing"

### Supplementary Methods

#### *Curation of gene networks*

We collected various network models for functional assessments of drug DEGs in PharmOmics. The networks included the STRING network (1) and tissue-specific gene regulatory networks. STRING is a composite network containing various types of interaction information (protein-protein interactions, gene co-expression, text mining, etc.) across different tissues. We filtered the network using a threshold of edge confidence score >= 95% to retain high confidence network nodes and edges. In addition to STRING, we also used tissue-specific gene regulatory networks, specifically Bayesian networks (BNs) of various tissues constructed using a previously established method (2,3) based on transcriptomic and genetic data from individual tissues (**Supplementary Table 11**). For each data set, 1,000 BNs with different random seeds were reconstructed using Monte Carlo Markov Chain simulation and the model with the best fit for each network was determined. In the resulting set of 1,000 networks, only edges appearing in over 30% of the networks were included in a consensus network. This practice has been found to produce experimentally supported regulatory relations between genes (2,3). For each tissue, the union of nodes and edges from BNs of multiple mouse or human studies were used as tissue-specific networks.

#### *Curation of drug signatures from CMAP, LINC1000, and CREEDS for comparison with PharmOmics*

To compare PharmOmics with other established drug signature platforms for drug repurposing, we downloaded signatures from L1000FWD(8) (http://amp.pharm.mssm.edu/l1000fwd/download_page) which were well annotated for matched drug signature overlapping comparison. For CREEDS (10) (http://amp.pharm.mssm.edu/CREEDS/) repositioning, their web-based enrichr (11) tool was used to query disease signatures to their DrugMatrix library and outputs based on "combined score" implemented by enrichr were used. Finally, CMAP repositioning test were completed through query to website directly (https://clue.io/) and rank based CMAP scoring were used. For CMAP and L1000 results which are based on in vitro cell lines, results from all cell lines were summarized to represent common usage of in vitro studies. For CREEDS results where in vivo study is available, only corresponding tissue were included for comparability with PharmOmics. We compared PharmOmics with the CREEDS, CMAP and L1000 at the regimens that showed the best performance in drug repurposing analysis.

#### *Assessing the relationship between drug DEGs and known drug targets*

To evaluate whether drug DEGs can capture known drug targets, we used three methods to assess the relationship between drug DEGs and known drug targets by checking whether the drug DEGs in PharmOmics i) contained known drug targets, ii) informed on known target pathways, or iii) are within the adjacent neighborhood of known drug targets in gene networks. Gene networks used here are the STRING network (1) and BNs described above. In order to address random connections between drug DEGs and known drug targets, permutation of drug targets using all human gene symbols (~26,000 genes) included in PharmOmics was conducted. For each random drug target pool, 1,000 permutations were performed to draw random genes matching the gene number of each drug DEG list. Overlaps between the randomly generated drug targets with DEGs for each drug in PharmOmics based on each of the three methods above were considered the null distribution. Finally, the background overlaps across all drugs were summed to conduct a Chi-square test to assess if drug DEGs can more accurately retrieve true drug targets than randomly generated drug targets.

***Curation of disease gene signatures for drug repositioning***

To test the potential of PharmOmics drug signatures for drug repositioning, we curated disease gene signatures for hyperlipidemia and non-alcoholic fatty liver disease (NAFLD). Hyperlipidemia was chosen as a test disease because numerous positive control drugs are available to assess the performance of PharmOmics in retrieving the known drugs compared to other existing drug repositioning tools. NAFLD was chosen as another test case since no effective drugs are currently available for this condition and our predictions may help guide future drug development.

The hyperlipidemia signatures were derived from two resources: i) genes and pathways identified by the MergeOmics pipeline (4) based on low-density lipoprotein cholesterol (LDL) genome-wide association study (GWAS) summary statistics data (12), and ii) genes based on mechanistic and therapeutic evidence collected by the Comparative Toxicogenomics Database (CTD) (13) under Mesh ID D006949. These two different resources represent disease gene signatures derived from either GWAS inference or literature-based system. NAFLD gene signatures were retrieved from i) studies of NAFLD mouse model (14) from a large systems genetics cohort comprised of hundreds of mice from ~100 genetically diverse strains, and ii) CTD gene signature under Mesh ID D065626. As additional test cases, we also retrieved gene signatures for chemical induced liver injury under CTD Mesh ID D056486, for acute kidney injury under CTD Mesh ID D058186, for hepatitis under CTD Mesh ID D006527, for hyperuricemia under CTD Mesh ID D033461 and for type 2 diabetes under CTD Mesh ID D003924.

***Measurement of similarity between signatures of drugs, ADRs and diseases***

We used two different methods to determine similarities between two signatures (e.g, a drug signature vs. a disease or ADR signature, or a drug signature vs. signature of another drug). The first method uses a signed Jaccard score based on upregulated genes from the first signature set (*a1*), upregulated genes from the second signature set (*b1*), downregulated genes from the first signature set (*a2*) and downregulated genes from the second signature set (*b2*). The Jaccard score was defined in the following formula:

$$J(A,B) = \frac{|A \cap B|}{|A \cup B|}$$

$$signed\ Jaccard\ score = J(a1,b1) + J(a2,b2) - J(a1,b2) - J(a2,b1)$$

If there is no direction from disease signature (*a)*, Jaccard score was determined based on simple overlapping of disease signature (*a)* and drug signatures without considering direction (*b1, b2*).

The second method to determine similarities between two signatures is based on a distance measure derived from the mean of shortest path lengths between network key drivers of a drug gene signature (A) and a disease signature (B) in a given Bayesian gene regulatory network (BN) based on a key driver analysis (see below for details). This distance measure is adapted from a previous study using protein interaction networks (15).

$$\mathrm{distance(B,A)} = \frac{1}{\|\mathrm{A}\|}\sum_{\mathrm{a \in A}} \mathrm{min\ b \in B\ distance(b,a)}$$

To reduce variations in result, only signatures with more than 10 genes were included in analysis. To obtain a null distribution for shortest path lengths, we permuted genes with the same degree as the drug/disease/ADR genes in each network 1,000 times and calculated a z-score based on the mean and standard error of the null distribution. ROCR package(16) were used to assessed performance of Jaccard score and network based z-score in disease repositioning.

**Supplementary Tables**

**Supplementary Table 1**. Distribution of treatment sample size in dose/time segregated dataset

| | | | | |
|---|---|---|---|---|
| Sample size | 1 | 2 | 3 | 4 |
| dataset number | 13 | 2491 | 9373 | 1 |
| percentage | 0.11% | 20.90% | 78.65% | 0.01% |
| Sample size | 5 | 6 | 7 | 9 |
| dataset number | 6 | 31 | 1 | 1 |
| percentage | 0.05% | 0.26% | 0.01% | 0.01% |

**Supplementary Table 2.** The capacity of drug signatures to retrieve known drug targets based on various methods.

*Only signatures with significant pathway annotations were included in the analysis.

| Method | | Median gene number count | Target detection rate | Target detection rate from randomized gene symbols | Statistical significance compared to random genes |
|---|---|---|---|---|---|
| gene overlap | PharmOmics dose/time segregated | 414 | 22.7% | 12.6% | $p < 0.001$ |
| | PharmOmics meta | 427 | 22.1% | 10.5% | $p < 0.001$ |
| | CREEDs | 387 | 14.1% | 7.1% | $p < 0.001$ |
| | L1000 | 450 | 17.2% | 7.4% | $p < 0.001$ |
| Pathway (GOBP and KEGG)* | PharmOmics dose/time segregated | 1816 | 59.1% | 23.5% | $p < 0.001$ |
| | PharmOmics meta | 1402 | 62.4% | 26.2% | $p < 0.001$ |
| Network (one-edge neighbors of DEGs in STRING network) | PharmOmics dose/time segregated | 2910 | 41.7% | 22.5% | $p < 0.001$ |
| | PharmOmics meta | 1143 | 41.8% | 17.4% | $p < 0.001$ |
| Network (one-edge neighbors of liver DEGs in liver BN) | PharmOmics dose/time segregated | 6673 | 60.2% | 38.1% | $p < 0.001$ |
| | PharmOmics meta | 4258 | 71.8% | 44.7% | $p < 0.001$ |
| Network (one-edge neighbors of liver DEGs in mismatched kidney BN) | PharmOmics dose/time segregated | 1863 | 47.0% | 24.3% | $p < 0.001$ |
| | PharmOmics meta | 861 | 40.5% | 19.2% | $p < 0.001$ |

**Supplementary Table 3. Comparison of drug repositioning performance between PharmOmics and other existing platforms for hyperlipidemia.** Drug pool for each database was limited to FDA approved drugs to match the drug selection criteria in PharmOmics to make results comparable. Significance were defined at the recommended cutoffs for each platform: z-score < -2.33 in PharmOmics, overlap BH adjusted p < 0.05 in CREEDS, L1000 and PharmOmics dose/time segregated Jaccard, and connection score > 95 or < -95 in CMAP query system. For CMAP and L1000, drug signatures from all cell lines (CMAP_all or L1000_all) or from the hyperlipidemia relevant liver cell line HepG2(CMAP_HEPG2 or L1000_HEPG2) were used.

| Drug signature platform | Total FDA listed drugs | Significant drugs (% total) | Known hyperlipidemia drugs | Significant hyperlipidemia drugs (% known drugs) | Balanced accuracy |
|---|---|---|---|---|---|
| PharmOmics meta_liver[*] | 263 | 54 (20.2%) | 10 | 6 (60%) | 69.6% |
| PharmOmics dose/time segregated_liver - network | 369 | 29 (7.9%) | 13 | 9 (69.2%) | 81.8% |
| PharmOmics dose/time segregated_liver - Jaccard | 369 | 171 (46.3%) | 13 | 12 (92.3%) | 73.8% |
| CMAP | 934 | 264 (28.6%) | 15 | 8 (53.3%) | 62.7% |
| CMAP_HEPG2 | 667 | 135 (20.3%) | 13 | 1 (7.7%) | 43.6% |
| L1000 | 867 | 428 (49.3%) | 14 | 8 (57.1%) | 53.9% |
| L1000_HEPG2 | 153 | 37 (24.2%) | 5 | 0 (0%) | 45.4% |
| CREEDS_liver | 281 | 257 (91.4%) | 12 | 12 (100%) | 54.4% |

[*]**Meta signature only uses drugs of platforms with similar genes assessed, which removed Codelink platform from drugMatrix database reducing the drug number.**

**Supplementary Table 4.** Prediction percentile of FDA approved gout treatment drug based on hyperuricemic signatures from CTD database across different platforms tested. HEPG2 results from both L1000 and CMAP were retrieved for tissue specificity comparison.

| Drugname | PharmOmics dose/time seg Network | PharmOmics dose/time seg Jaccard | CREEDS | CMAP_ HEPG2 | CMAP | L1000_ HEPG2 | L1000 |
|---|---|---|---|---|---|---|---|
| Allopurinol | 0.846 | 0.317 | NA | NA | NA | NA | NA |
| Benzbromarone | 0.946 | 0.451 | NA | 0.313 | 0.475 | 0.408 | 0.895 |
| Colchicine | 0.889 | 0.271 | NA | NA | NA | NA | NA |
| Febuxostat | NA | NA | NA | 0.940 | 0.821 | NA | NA |
| Probenecid | NA | NA | NA | 0.043 | 0.211 | NA | 0.176 |
| Sulfinpyrazone | NA | NA | NA | NA | 0.524 | NA | 0.176 |
| **Median** | **0.889** | **0.317** | **NA** | **0.313** | **0.499** | **0.408** | **0.176** |
| **Mean** | **0.893** | **0.346** | **NA** | **0.432** | **0.508** | **0.408** | **0.416** |

**Supplementary Table 5.** Prediction percentile of non-steroid anti-inflammatory drugs based on hepatitis signatures from CTD database across different platforms tested. HEPG2 results from both L1000 and CMAP were retrieved for tissue specificity comparison.

| **Drugname** | **PharmOmics dose/time seg Network** | **PharmOmics dose/time seg Jaccard** | **CREEDS** | **CMAP _HEPG 2** | **CMAP** | **L1000_ HEPG2** | **L1000** |
|---|---|---|---|---|---|---|---|
| Aspirin | 0.870 | 0.874 | 0.856 | NA | 0.102 | NA | NA |
| Bromfenac | 0.618 | 0.774 | 0.705 | NA | 0.777 | NA | NA |
| Diclofenac | 0.593 | 0.821 | 0.521 | 0.204 | 0.503 | NA | 0.645 |
| Ibuprofen | 1.000 | 0.898 | 0.972 | NA | 0.390 | NA | 0.436 |
| Indomethaci n | 0.913 | 0.753 | 0.989 | NA | NA | NA | NA |
| Ketorolac | 0.780 | 0.889 | 0.993 | 0.438 | 0.791 | NA | 0.309 |
| Mefenamic acid | 0.924 | 0.694 | NA | NA | NA | NA | NA |
| Naproxen | 0.569 | 0.472 | 0.594 | 0.753 | 0.937 | NA | 0.340 |
| Phenylbutaz one | 0.976 | 0.967 | NA | 0.066 | 0.592 | NA | 0.185 |
| Sulindac | 0.932 | 0.724 | 0.872 | NA | 0.286 | NA | 0.386 |
| Acemetacin | NA | NA | 0.270 | NA | NA | NA | NA |
| Benzydamin e | NA | NA | NA | 0.120 | 0.484 | NA | 0.453 |
| Dexketoprof en | NA | NA | NA | 0.256 | 0.327 | NA | 0.231 |
| Diflunisal | NA | NA | NA | 0.781 | 0.350 | NA | 0.720 |
| Fenbufen | NA | NA | NA | 0.889 | 0.353 | NA | 0.015 |
| Ketoprofen | NA | NA | NA | 0.076 | 0.808 | NA | 0.015 |
| Oxaprozin | NA | NA | NA | 0.633 | 0.418 | NA | 0.179 |
| Piroxicam | NA | NA | NA | 0.034 | 0.898 | NA | 0.851 |
| Tenoxicam | NA | NA | NA | 0.694 | 0.340 | NA | 0.238 |
| Tolmetin | NA | NA | NA | 0.870 | 0.273 | NA | 0.138 |
| Ampiroxica m | NA | NA | NA | NA | 0.203 | NA | 0.043 |
| Loxoprofen | NA | NA | NA | NA | 0.911 | NA | 0.392 |
| Mofezolac | NA | NA | NA | NA | 0.854 | NA | 0.330 |
| Nabumetone | NA | NA | NA | NA | 0.685 | NA | 0.309 |
| Flurbiprofen | NA | NA | NA | NA | NA | NA | 0.818 |
| **Median** | **0.892** | **0.797** | **0.856** | **0.438** | **0.484** | **NA** | **0.320** |
| **Mean** | **0.818** | **0.787** | **0.752** | **0.447** | **0.537** | **NA** | **0.352** |

**Supplementary Table 6.** Prediction percentile of FDA approved anti-diabetic drug based on type2 diabetes signatures from CTD database across different platforms tested. HEPG2 results from both L1000 and CMAP were retrieved for tissue specificity comparison.

| Drugname | PharmOmics dose/time seg Network | PharmOmics dose/time seg Jaccard | CREEDS | CMAP_ HEPG2 | CMAP | L1000_ HEPG2 | L1000 |
|---|---|---|---|---|---|---|---|
| **Sulfonylurea drugs** | | | | | | | |
| Chlorpropamide | 0.547 | 0.279 | NA | NA | NA | NA | NA |
| Glimepiride | 0.241 | 0.450 | 0.456 | 0.835 | 0.343 | NA | 0.717 |
| Glipizide | 0.260 | 0.301 | 0.523 | 0.498 | 0.845 | NA | 0.005 |
| Nateglinide | 0.179 | 0.480 | 0.231 | 0.289 | 0.590 | NA | 0.560 |
| Tolazamide | 0.046 | 0.397 | 0.267 | 0.465 | 0.120 | NA | 0.770 |
| Tolbutamide | 0.699 | 0.136 | NA | 0.832 | 0.290 | NA | 0.361 |
| Gliquidone | NA | NA | NA | 0.940 | 0.678 | NA | 0.270 |
| Repaglinide | NA | NA | NA | NA | 0.779 | NA | 0.376 |
| **Median** | **0.251** | **0.349** | **0.361** | **0.665** | **0.590** | **NA** | **0.376** |
| **Mean** | **0.329** | **0.340** | **0.369** | **0.643** | **0.521** | **NA** | **0.437** |
| **PPAR gamma agonist** | | | | | | | |
| Pioglitazone | 0.688 | 0.298 | 0.854 | 0.906 | 0.656 | 0.350 | 0.826 |
| Rosiglitazone | 0.810 | 0.734 | 0.765 | 0.955 | 0.970 | NA | 0.680 |
| Troglitazone | 0.873 | 0.419 | 0.655 | 0.684 | 0.256 | NA | 0.976 |
| Ciglitazone | NA | NA | NA | 0.499 | 0.925 | NA | 0.023 |
| **Median** | **0.810** | **0.419** | **0.765** | **0.795** | **0.791** | **0.350** | **0.753** |
| **Mean** | **0.790** | **0.484** | **0.758** | **0.761** | **0.702** | **0.350** | **0.626** |

**Supplementary Table 11.** Data resources and references for the construction of Bayesian gene-gene regulatory networks.

| Tissue | Species | Dataset |
|---|---|---|
| Liver | Mouse | C57BL/6J x A/J mouse cross (17) |
| | | C57BL/6J x C3H ApoE -/- mouse cross (18,19) |
| | | C57BL/6J x C3H wildtype mouse cross (20) |
| | | C57BL/6J x BTBR Lepob mouse cross (21) |
| | Human | 427 individuals (22) |
| Kidney | Mouse | C57BL/6J x A/J mouse cross (17) |
| | | CAST, PWK and WSB mouse cross (23) |
| | | 47 strains of the collaborative mouse cross (24) |

**The following Supplementary Tables are uploaded as separate excel files.**

**Supplementary Table 7.** Network repositioning result for non-alcoholic fatty liver disease based on genetic pathways obtained from studies of female and male mice.

**Supplementary Table 8.** Submodule repositioning result based on signatures from CTD chemical induced liver injury

**Supplementary Table 9.** Cross-tissue comparison of Atorvastatin Pathways.

**Supplementary Table 10.** Cross-species comparison of Rosuvastatin Pathways.

### Supplementary Figures

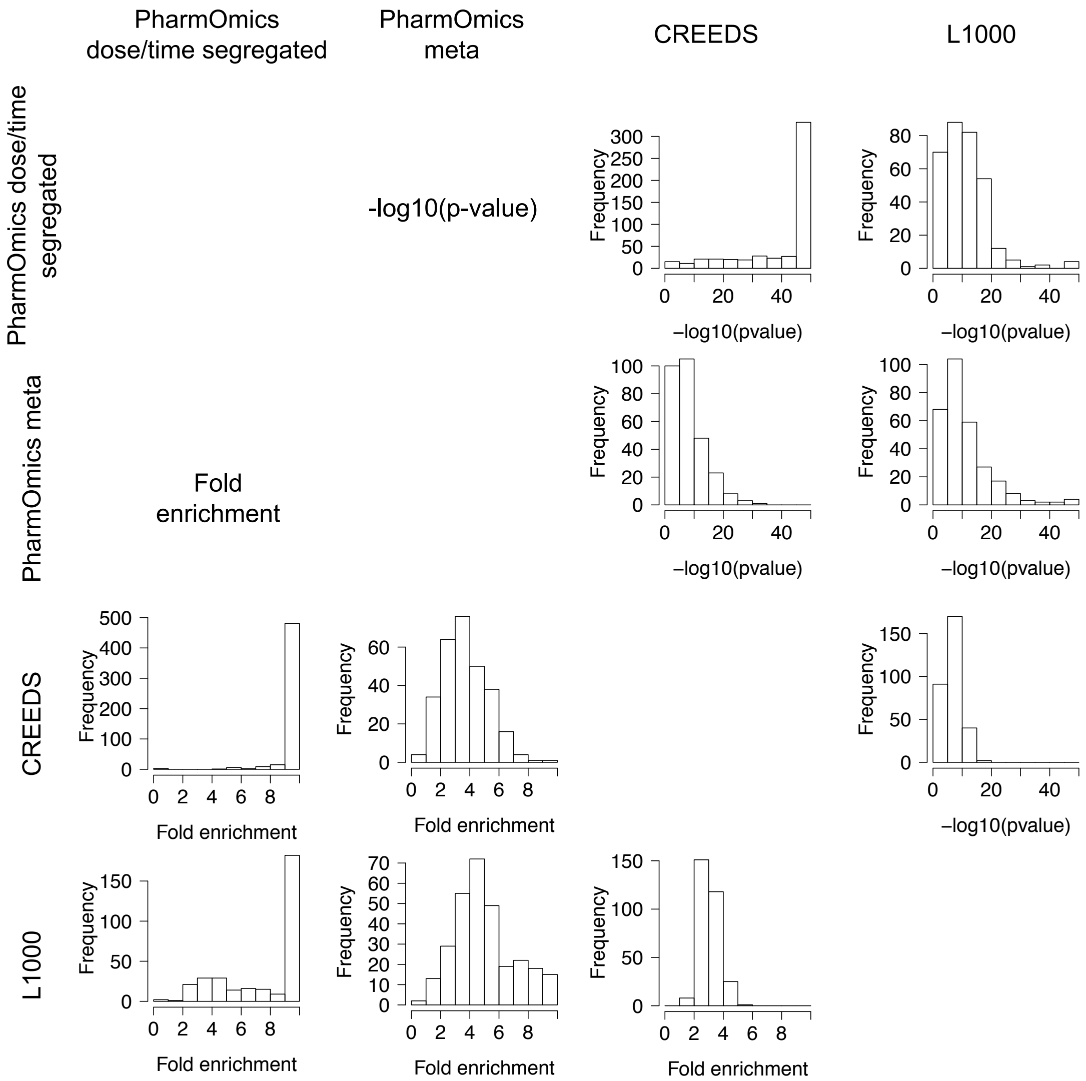


**Supplementary Figure 1. Comparison of drug signatures between PharmOmics and existing drug signature databases CREEDS and L1000**. For each drug, only the overlap scores for the best matched signatures between two databases are used. Lower left triangular matrix represents the histogram of the overlap fold enrichment scores calculated using hypergeometric test, and the upper right triangular matrix represents the histogram of the -log10(pvalue) from Fisher's exact test.

**Supplementary Figure 2.**

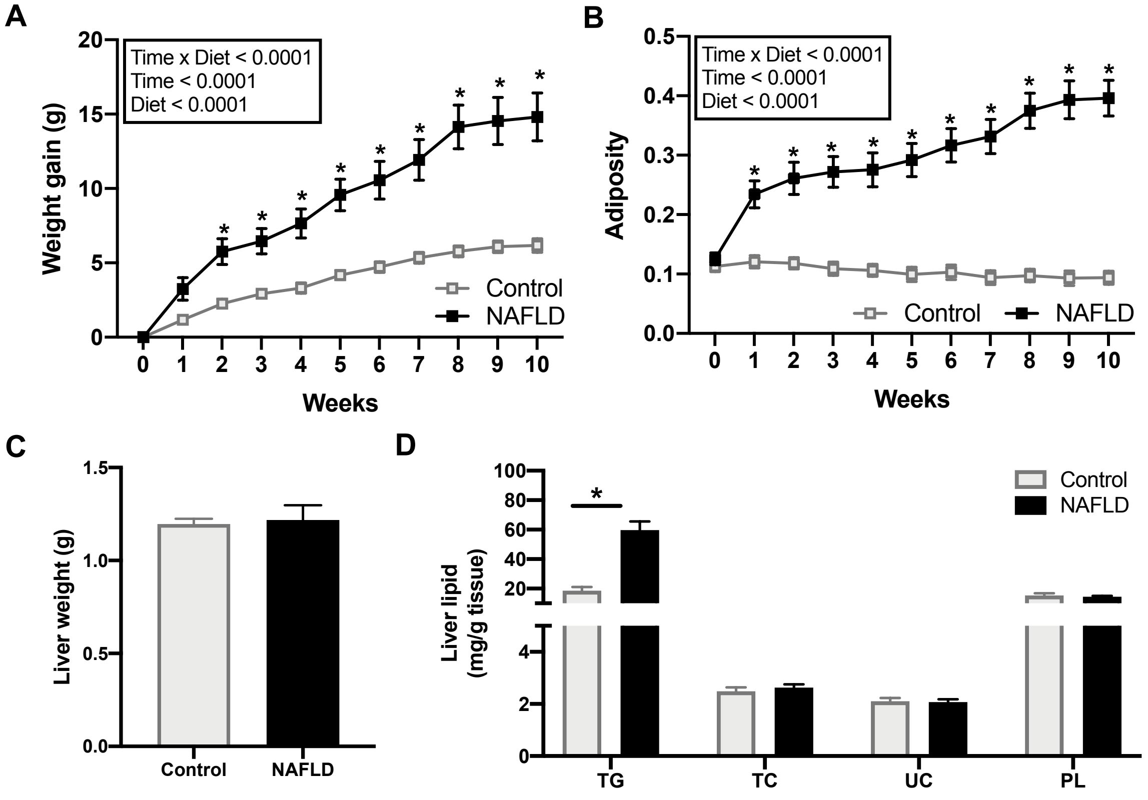


**Supplementary Figure 2. Effects of high fat high sucrose on body composition and liver lipids in C57BL/6J mice.** Time course of body weight gain of mice on control and high fat high sucrose diet for 10 weeks (A). Time course of adiposity of mice on control and high fat high sucrose diet for 10 weeks (B). Data was analyzed by two-way ANOVA followed by Sidak post-hoc analysis to examine treatment effects at individual time points (A and B). Bar plot of liver weight in mice on a control and high fat high sucrose diet (C). Bar plot of hepatic lipid levels in mice on a control and high fat high sucrose diet (D). Triglyceride (TG), Total Cholesterol (TC), Unesterified Cholesterol (UC), Phospholipid (PL). Data was analyzed using two-sided Student's t-test (C and D). P value <0.05 was considered significant and is denoted by an asterisk (*). Sample size n = 7-9/group. Control diet (Control); high fat high sucrose diet (NAFLD).

**Supplementary Figure 3.**

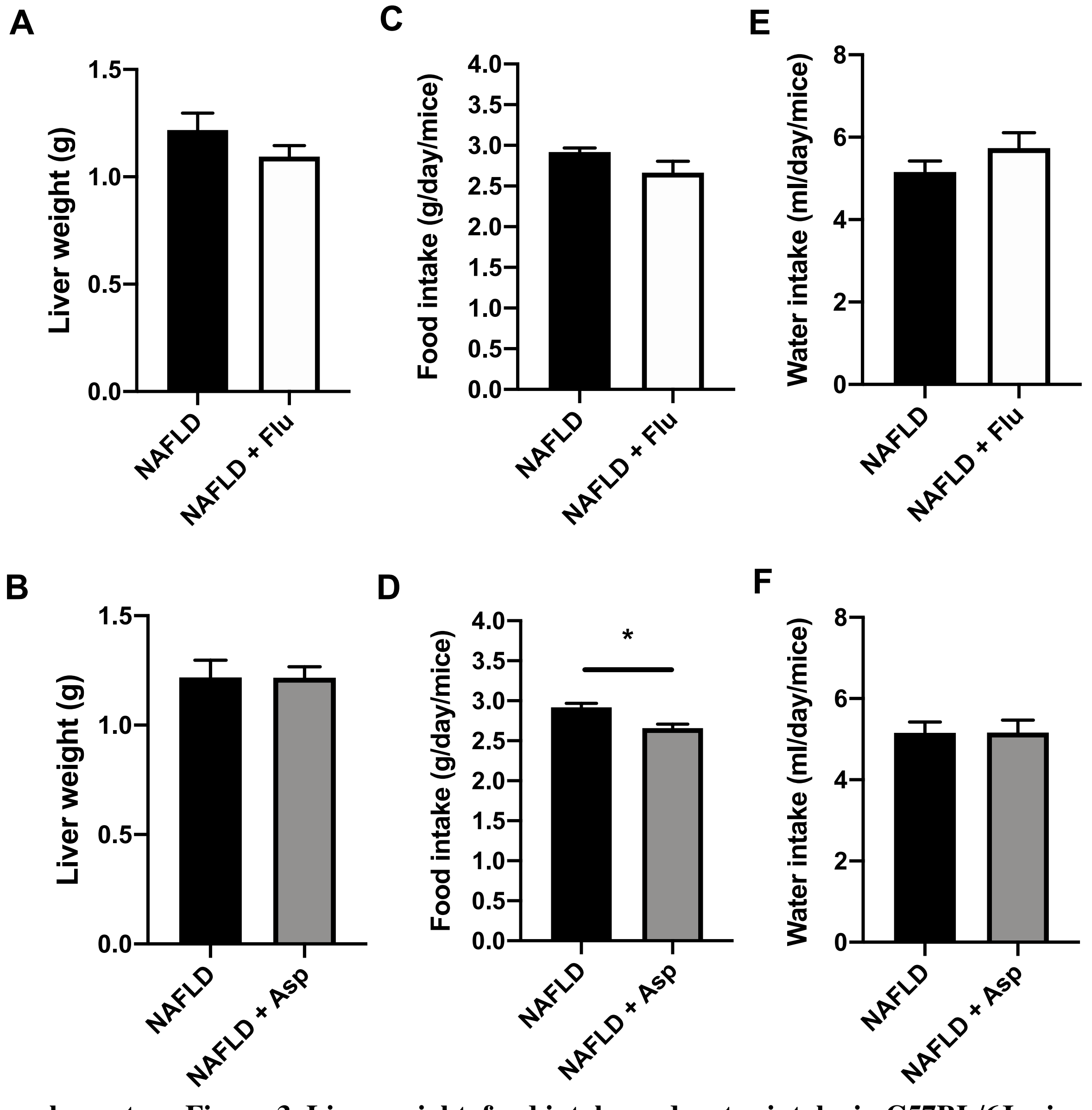


**Supplementary Figure 3. Liver weight, food intake and water intake in C57BL/6J mice.** Bar plot of liver tissue weight in mice on a high fat high sucrose diet with and without fluvastatin (A) or aspirin (B). Bar plot of food intake in mice on a high fat high sucrose diet with and without fluvastatin (C) or aspirin (D). Bar plot of water intake in mice on a high fat high sucrose diet with and without fluvastatin (E) or aspirin (F). Data was analyzed using two-sided Student's t-test. P value <0.05 was considered significant and is denoted by an asterisk (*). Sample size n = 7-9/group. High fat high sucrose group (NAFLD); High fat high sucrose with Fluvastatin (NAFLD + Flu); High fat high sucrose with Aspirin (NAFLD + Asp).

**Supplementary Figure 4.**

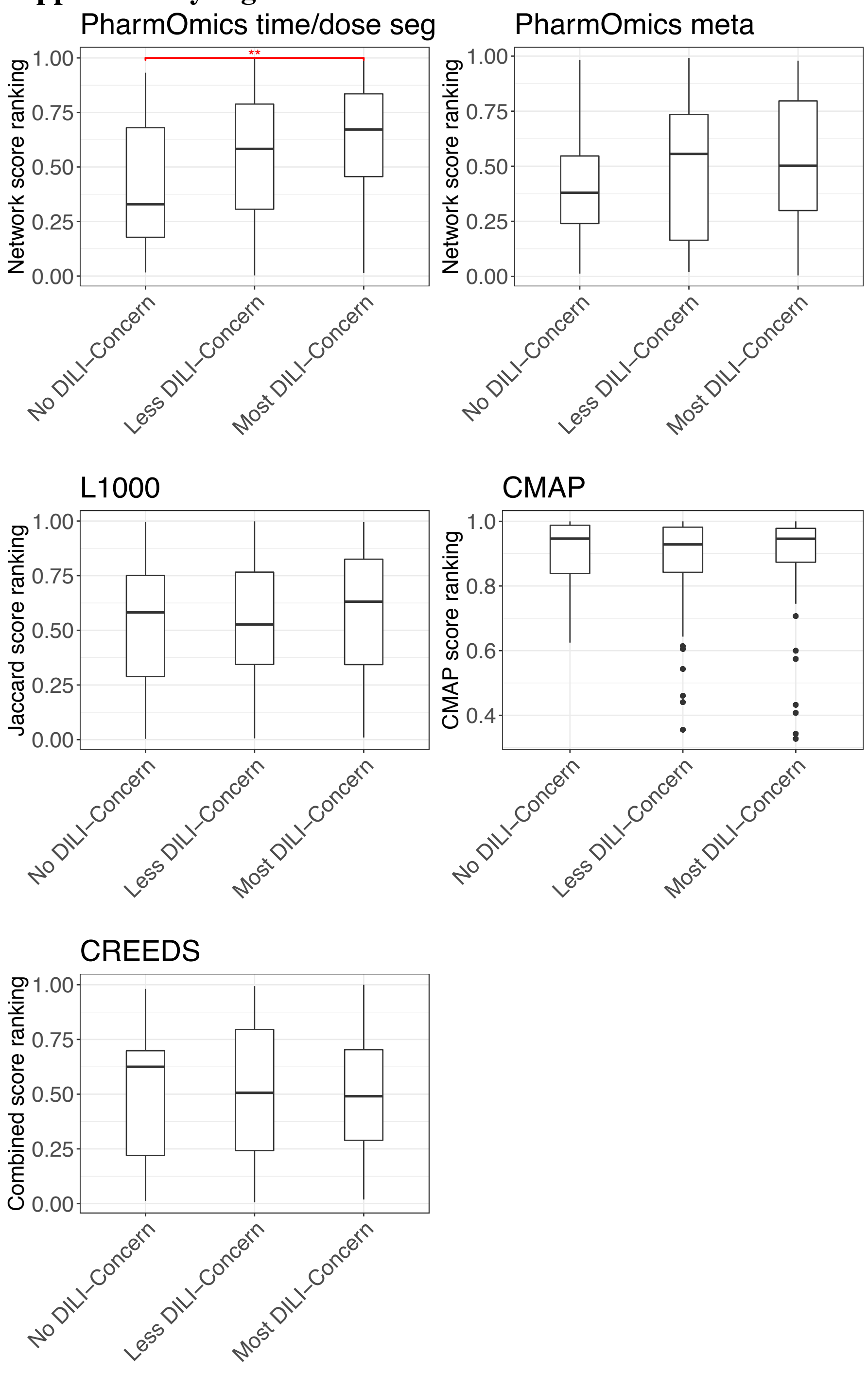


**Supplementary Figure 4.** Distribution of repositioning score (PharmOmics: network score ranking, CMAP: CMAP scores, L1000 and CREEDS: Jaccard score ranking) in three different DILI drug categories by repositioning of chemical induced liver injury score. For drugs with multiple dose and time points, only best score were used. ANOVA test with Tukey procedure were used for statistics. *, ** indicates $p < 0.0\ 5$ and $p < 0.01$ respectively. Boxplot interquartile range were defined as $25^{th}$ to $75^{th}$ percentile, median value were used for line inside the box. Upper and lower bounds were defined as 75th percentile + 1.5*IQR and 25th percentile – 1.5*IQR respectively.
