## Supplementary figures and images for "PharmOmics: A Species- and Tissue-specific Drug Signature Database and Online Tool for Drug Repurposing"

### Supplementary Figure 1

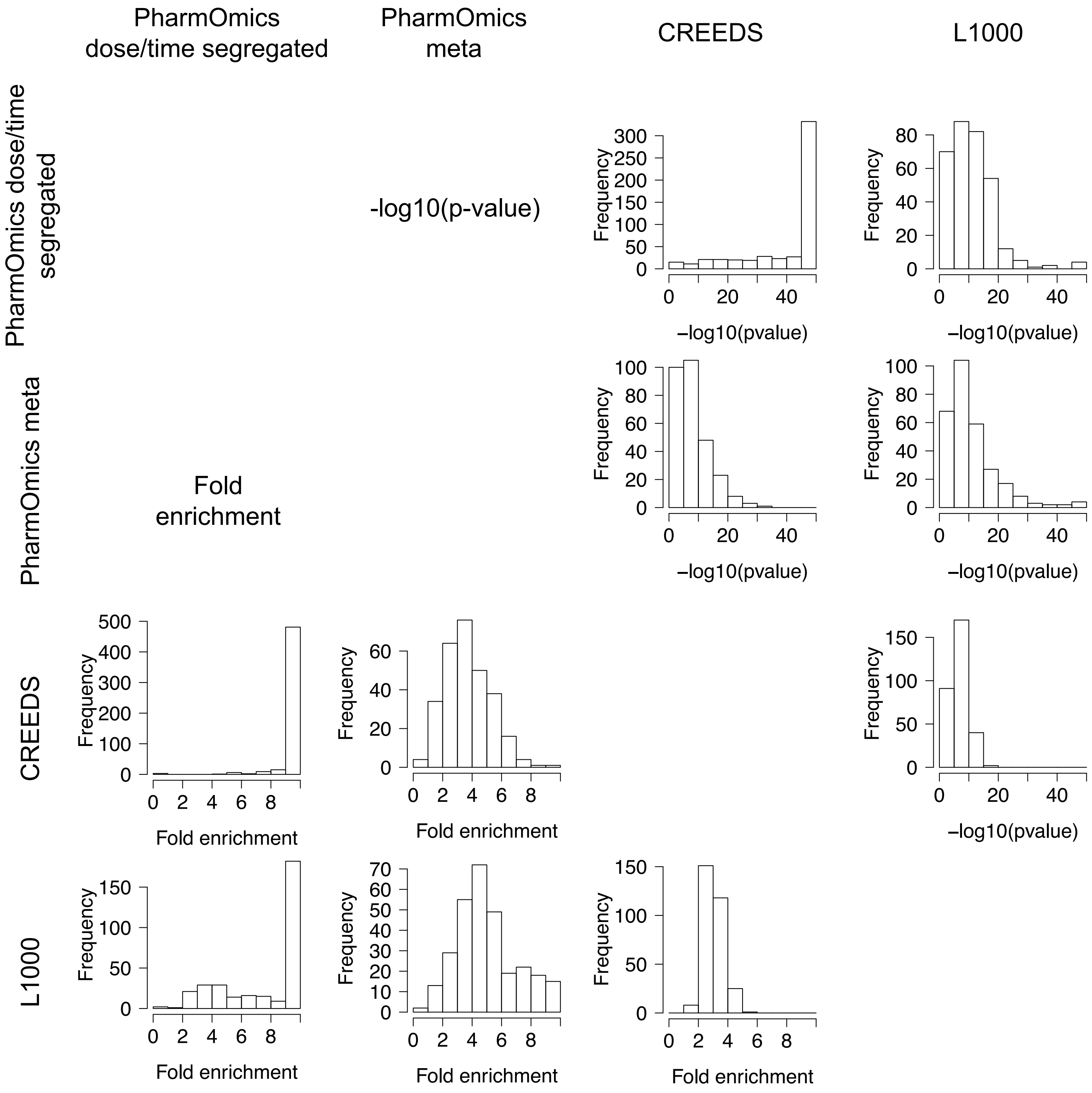

### Supplementary Figure 2

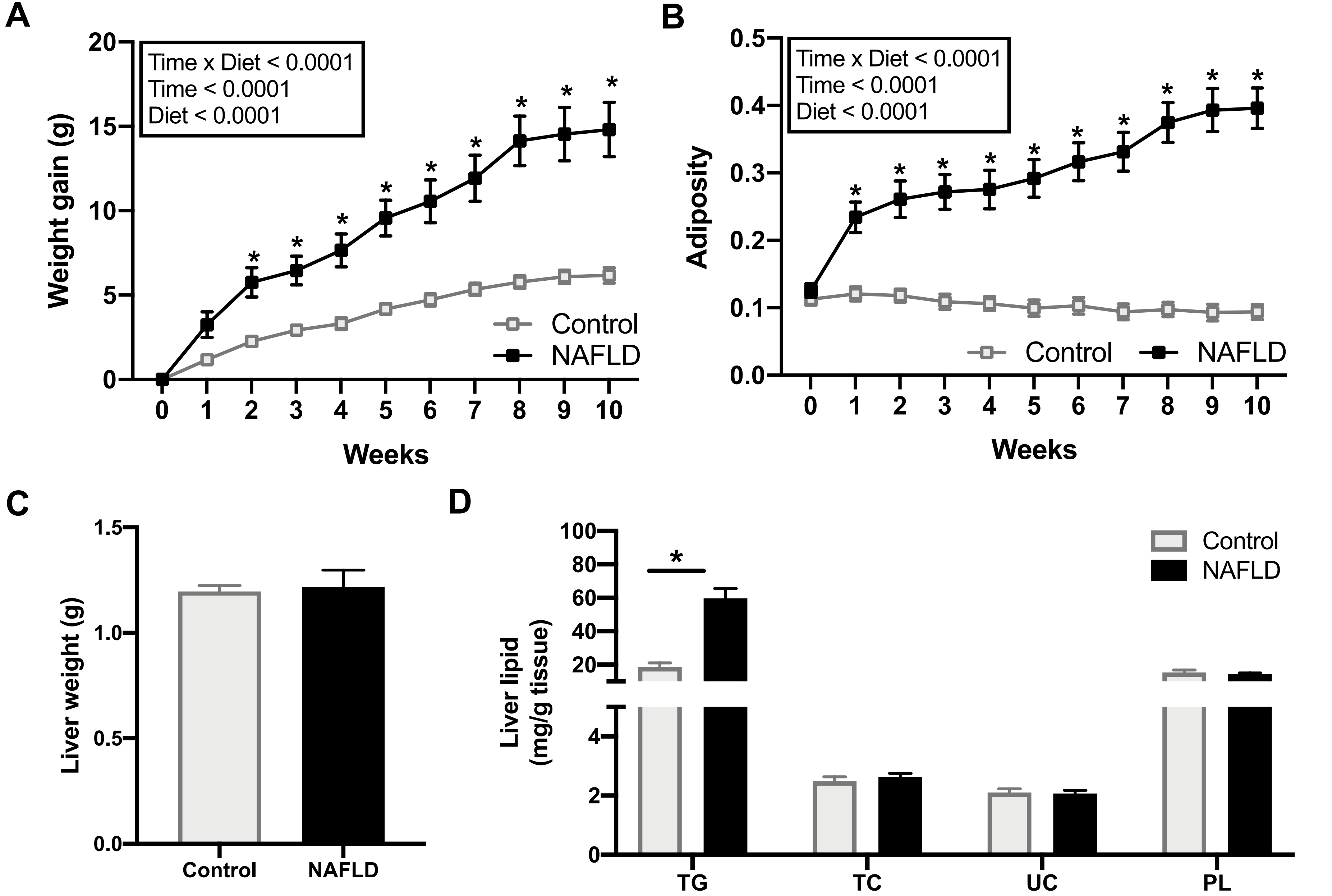

### Supplementary Figure 3

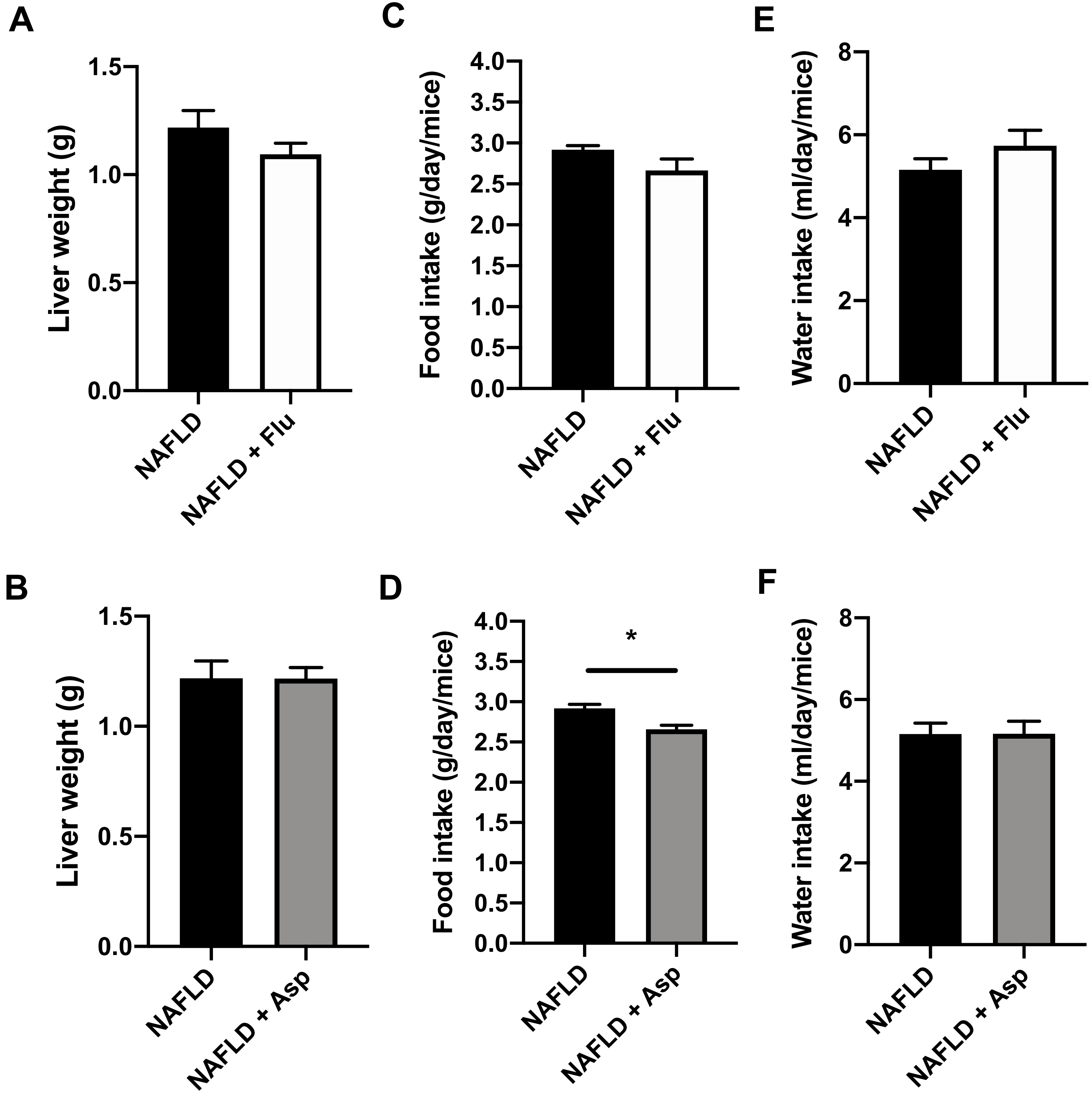

### Supplementary Figure 4

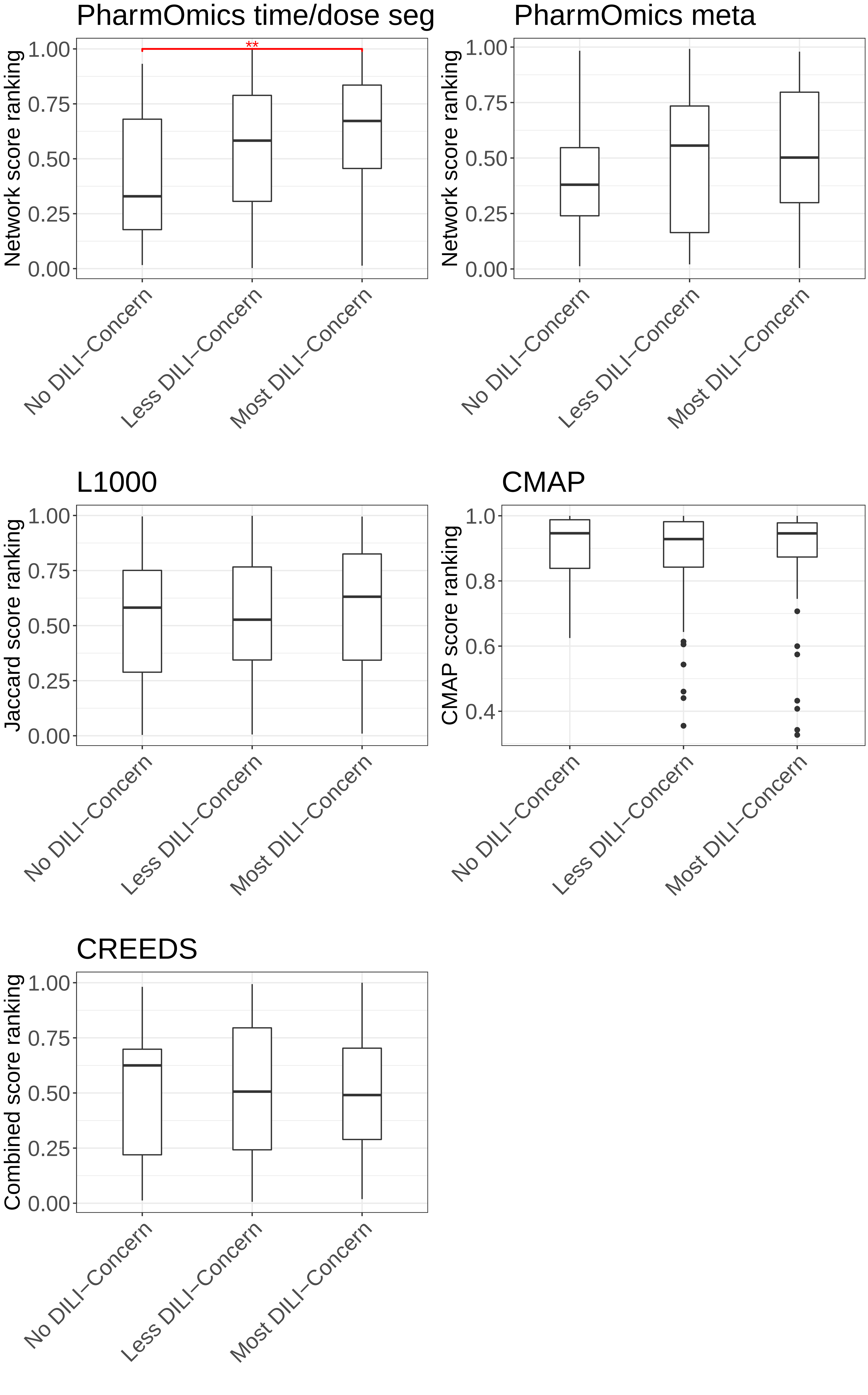
